# Microelectrode arrays enable directional stereo-EEG during kainate-mediated seizures

**DOI:** 10.64898/2026.06.11.731746

**Authors:** Ryan Shores, Takfarinas Medani, Anand Joshi, Camryn Matthews, Yash S. Vakilna, Jay Gavvala, Richard M. Leahy, Sandipan Pati, John C. Mosher, Nitin Tandon, John P. Seymour

**Author notes:** Correspondence to: John P. Seymour, Full address: 6500 Main Street, BRC Room 860F, Houston TX 77030.

## Abstract

Surgical planning for drug-resistant epilepsy often relies on stereo-EEG (sEEG) recordings obtained with cylindrical ring electrodes. Prior modeling studies, including lead-field analysis, suggest that microelectrodes distributed around an sEEG-sized insulating body offer superior source amplification and directional sensitivity that are not available with either a ring design or microelectrodes on micro-structures, e.g. Neuropixel. However, these advantages have not been demonstrated in seizure models.

This study evaluated high-density sEEG recordings using directional microelectrode arrays in a kainate-mediated rat model (n=6). Two 64-channel microelectrode arrays were implanted near the hippocampus, and the signals were spatially averaged to emulate virtual ring electrodes for comparison. Device locations were reconstructed and placed in copies of the Waxholm Space Rat Brain Atlas registered to subject-specific MRI scans.

In subjects exhibiting seizures (n=4), automated line length detection showed that microelectrode signals identified epileptiform activity sooner and with greater specificity than ring electrodes. In subjects that only seized post-kainate injection (n=3), manual review by a board-certified epileptologist confirmed that microelectrodes provided the earliest onset detection times. Furthermore, the microelectrode arrays’ high resolution revealed distinct instances of hyperactivity occurring at similar depths but from different directions — a feature indistinguishable to standard ring electrodes.

These results demonstrate that microelectrodes on a large insulating body significantly enhance signal quality and spatial localization. This technology offers a potential advancement over current clinical standards for identifying seizure foci during surgical planning.

**SIGNIFICANCE:** Surgical treatment for drug-resistant epilepsy depends on accurately identifying the brain regions where seizures begin. Current stereo-EEG electrodes sample activity with ring contacts that average signals around the probe shaft, potentially obscuring directional differences in nearby neural activity. This study shows that microelectrodes distributed around an sEEG-sized insulating body can improve seizure-related signal detection and spatial localization compared with ring-like recordings from the same implant locations. By resolving activity that conventional ring electrodes cannot distinguish, high-density directional sEEG may provide more informative recordings for seizure mapping. These findings support the development of next-generation intracranial electrodes for improving epilepsy surgical planning.

## Introduction

Epilepsy affects approximately 50 million people worldwide, and temporal lobe epilepsy is the most common focal epilepsy syndrome in adults (Téllez-Zenteno and Hernández-Ronquillo, 2012; World Health Organization, 2024). Although multiple antiseizure medications are currently available, more than one-third of patients remain drug resistant (Kwan and Brodie, 2000). For carefully selected patients with drug-resistant temporal lobe epilepsy, particularly those with hippocampal sclerosis, resective surgery can substantially improve remission rates and may achieve long-term seizure freedom in a large proportion of cases (Wiebe *et al*., 2001; Jayalakshmi *et al*., 2023). Although focal-onset epilepsies account for roughly 60% of adult epilepsy cases, identifying the true seizure-onset network remains challenging (Téllez-Zenteno and Hernández-Ronquillo, 2012).

Epileptic seizures arise from abnormal neuronal networks characterized by excessive excitability and synchrony (Ren *et al*., 2021; World Health Organization, 2024). Because neural tissue is extensively interconnected, these networks can extend across multiple subregions, spreading seizure activity into areas that are not themselves critical for seizure generation (Chowdhury *et al*., 2021; Ren *et al*., 2021). As a result, diagnostic tools must distinguish truly epileptogenic tissue from surrounding irritative and otherwise normally functioning tissue. Standard presurgical evaluation begins with noninvasive electroencephalography (EEG) to approximate the cortical regions relevant to the patient’s epileptogenic zone. However, scalp EEG has limited spatial resolution because signals are distorted by the scalp, skull, cerebrospinal fluid, and intersubject differences in tissue conductivity (van Mierlo *et al*., 2020). When lesion imaging and scalp EEG are inconclusive, invasive intracranial recordings are often required.

The two principal modalities are subdural electrode arrays and depth electrode arrays. Subdural arrays provide dense cortical surface sampling but require craniotomy and are limited in their ability to sample deep structures (Jehi *et al*., 2021; Wu *et al*., 2024). By contrast, stereoelectroencephalography (sEEG) with depth electrodes better samples deep and bilateral networks and is generally associated with lower overall complication and infection rates than subdural monitoring. However, conventional cylindrical contacts remain relatively coarse and nondirectional, so localization often depends on integrating information across multiple contacts and trajectories (Jehi *et al*., 2021; Abrego *et al*., 2023; Wu *et al*., 2024).

The epileptogenic and irritative zones represent unknown tissue volumes, yet the clinical team must identify the epileptogenic target for resection, ablation, or neuromodulation. Among available interventions, resection generally offers the highest probability of sustained seizure freedom, but it also carries the greatest risk of neurological or cognitive deficit when functional tissue is removed (Vakharia *et al*., 2018; Yan and Ibrahim, 2019). After language-dominant temporal lobe resection, postoperative naming decline occurs in approximately one-third of patients (Sherman *et al*., 2011). Likewise, when seizure onset lies near eloquent cortex, surgery carries meaningful risk of motor or language deficits, making precise functional mapping and margin planning essential (Vakharia *et al*., 2018; Yan and Ibrahim, 2019). Therefore, improved source localization may help maximize seizure freedom while limiting functional morbidity, particularly when focal ablation rather than a larger resection is chosen.

This study compares microelectrode arrays with conventional ring contacts in a rodent seizure model. The <u>di</u>rectional and <u>sc</u>alable (DiSc) depth arrays evaluated here retain a clinically compatible 0.8mm form factor while providing directional sensitivity and field potential amplification (Abrego *et al*., 2023; Willis *et al*., 2026). In humans, microseizures have been demonstrated with hybrid micro/macro subdural arrays (Stead *et al*., 2010)] and with high-density micro-electrocorticography arrays (Sun *et al*., 2022). While both theory and early empirical evidence support the claim that microelectrodes on a mesoscale insulating body can increase signal amplitude and localization, the epileptogenic sources and potential noise from microcontacts (Stacey *et al*., 2012) requires more investigation. Here, we propose using DiSc arrays to record local field potentials and microseizures in a rodent seizure model induced by intra-amygdaloid administration of the glutamate agonist kainic acid (KA). In this model, epileptiform activity propagates from the amygdala to the hippocampus, where seizure-associated neuronal injury has been well documented (Ben-Ari, Tremblay and Ottersen, 1980; Rusina, Bernard and Williamson, 2021). Accordingly, we implanted two 64-channel DiSc arrays into the hippocampus of Sprague Dawley rats and recorded before and after KA administration into the basolateral amygdala (BLA). We further demonstrate how Brainstorm, together with the Brainstorm-DUNEuro finite-element modeling pipeline, can be used to generate high-resolution source localization of the recorded activity (Tadel *et al*., 2011; Medani *et al*., 2023). This study offers a novel approach for investigating the spatial organization of seizure networks in vivo.

## Materials and methods

### Experimental Design and Statistical Analysis

Six male Sprague Dawley rats were included in this experiment. Comparison of device configurations was originally modeled as a repeated measures ANOVA using within subject factors with one group and 12 measurements – three device configurations with four 1.3mm spans per device. Assuming an error probability of 0.05, power of 0.8, correlation of 0.7 and nonsphericity correction of 1, power analysis revealed that at least 8 devices would be required for sufficient power. With two devices implanted per subject, this is reduced to a minimum of 4 subjects. Power analysis was done with GPower. All other statistics were computed using Python. The t-tests were performed with the “scipy” Python package. Generalized Estimating Equations and Linear Mixed Models were implemented with the “statsmodels.api” Python library. The Cox proportional hazard model was fit to the time-to-event latencies using the “lifelines” Python package.

### Acute Kainic Acid Surgery

All procedures were conducted in accordance with regulations established by the Rice University Institutional Animal Care and Use Committee. Male Sprague Dawley Rats were acquired from Charles River and housed for at least a week prior to use in experiments. All in vivo animal work done with a given subject was performed in a single day (Figure 1-A).

**Figure 1.**
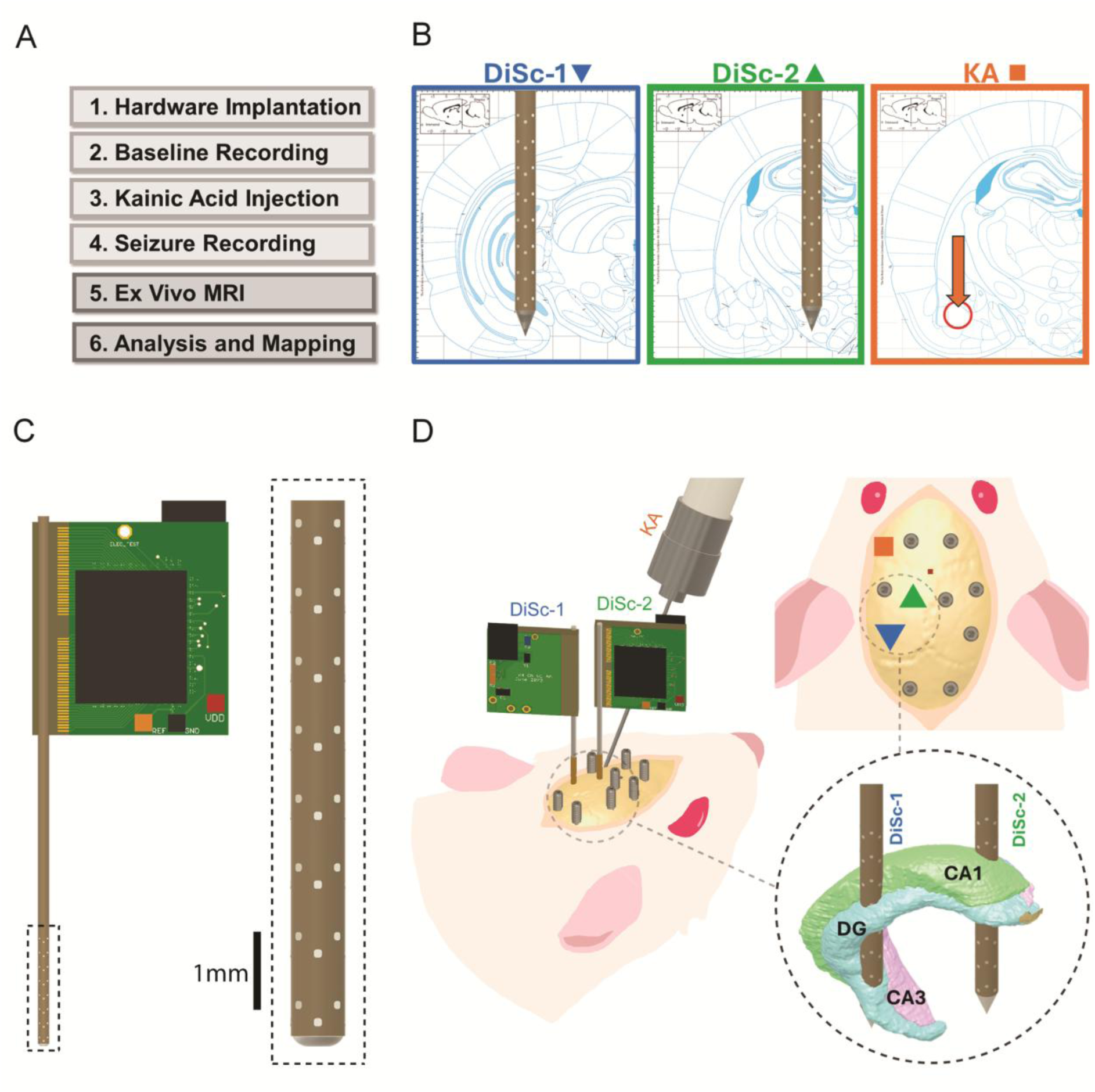
Acute Seizure Timeline and Hardware Layout. (**A**) Experimental timeline. Hardware implantation and recording sessions were carried out in a single day. Ex vivo MRI and data analysis were performed afterwards. (**B**) Coronal atlas slices at DiSc array implantation targets overlaid on a simplified image from (Paxinos and Watson, 2007) at: AP: -6.12 (left), AP: -2.76 (center), and AP: -2.52 (right). Background gridlines are 1mm apart. (C) CAD model of a DiSc 64-channel microelectrode array. Of the 128 electrode contacts on the array, only 64 are active due to amplifier constraints; only the active electrodes are shown. Microelectrodes are arranged in 8 rows and 8 columns with a total span of 7.5 mm. (D) Hardware layout on the skull shown from perspective (left) and top-down view (upper right). The intracranial arrays flank the left hippocampus from either side (lower right). The layouts are depicted to scale.

### Animal Preparation

On the day of surgery, rats were transported from their home cages to the surgical suite and weighed. The subjects were then induced under 4% isoflurane with room air using a SomnoSuite micro-anesthesia unit. Once breathing slowed and subjects proved unresponsive to a foot pinch, they were removed from the induction chamber and mounted to a dual arm stereotactic frame on top of a warm water heat therapy pad. The rest of the procedure was carried out under 1.5-2.5% isoflurane.

### Surgical Site Preparation

Once mounted to the stereotaxic bite bar, 5% topical lidocaine was applied to ear bars which were then used to stabilize the head. An injection of Ethiqa XR extended-release buprenorphine (0.65mg/kg) was then administered subcutaneously for multimodal analgesia with general anesthesia. The fur on the subject’s scalp was shaved and then the surgical site was cleaned with alternating administrations of betadine and ethanol. Ophthalmic ointment was also applied to the eyes. Vitals were monitored using a rectal thermometer coated in petroleum jelly and a pulse oximeter clamped to the foot.

An incision was made to expose the cranial sutures then the head was leveled such that bregma and lambda occupied the same DV coordinate within an error of (+/- 0.1mm). A microinjection syringe was subsequently mounted to the stereotaxic frame at a 30-degree pitch from the standard vertical position and loaded with KA diluted in 1x PBS (2.67 ng/nL). The needle tip was zeroed at bregma before being moved to the extremities of the range of the stereotaxic arm to prevent the syringe body from obstructing the surgical field (Figure 1-D).

A custom 3D-printed probe holder was mounted on the second arm of the stereotaxic frame. An 800um diameter stainless-steel rod was then loaded into the probe holder using a standard probe holder corner clamp such that the rods extended below the bottom of the tool. A burr hole was then drilled with a 1.4mm dental burr to serve as the entry point of the KA syringe.

### Surface Electrode Implantation

Three different styles of surface electrodes were implanted in the animals used in this study – electrocorticography (ECoG) with stainless steel bone screws, ECoG with microwires, and sub-scalp EEG with a 32-channel, thin-film electrode array. Each subject received only one style of surface electrode in addition to the intracortical DiSc arrays (Table 1). The microwire ECoG method is not recommended but has been included since Subject 5 received this hardware.

**Table 1.** Acute Seizure Surgery Details.

| **Subject** | **Analysis Group** | **Seizure Onsets After KA Treatment** | **Spiking in Baseline** | **Surface Hardware** | **Implantation Depth (DV)** | **Guide Wire Diameter** | **KA Injected** |
| --- | --- | --- | --- | --- | --- | --- | --- |
| **1** | 1 | 3 | No | ECoG (screw) | -8.5mm | 500um SS, 700um PTFE | 900nL |
| **2** | 1 | 3 | No | ECoG (screw) | -8.5mm | 500um SS, 700um PTFE | 600nL |
| **3** | 1 | 1 | No | Sub-scalp EEG | -8.5mm | Tapered 800um SS | 300nL |
| **4** | 2 | 1 | Yes | ECoG (screw) | -10.7mm | 500um SS, 700um PTFE | 300nL |
| **5** | 3 | 0 | Yes | ECoG (wire) | -8.5mm | 500um SS, 700um PTFE | 900nL |
| **6** | 3 | 0 | No | Sub-scalp EEG | -9.7mm | Tapered 800um SS | 600nL |

### Surface Electrode Implantation Method A: Bone Screw ECoG

This was the primary surface recording method (n=3). Markings for 7 ECoG bone screw electrodes were made with an additional mark at to denote the reference electrode. The exact locations have been provided as supplemental text in the appendix (Supplemental Item 1). Pilot holes were drilled at each implantation site with a 1.35mm stainless steel drill bit. Prior to surgery, stainless-steel, size 0-80 cup point set screws were soldered to 30-gauge silver plated copper wire with a metal pin on the opposite end. Once all the coordinates for the implanted hardware were marked, each pre-soldered lead was driven into its respective hole with a hex driver (Figure 1-D).

### Surface Electrode Implantation Method B: Thin-film Skull EEG

This was the secondary surface recording method (n=2). A custom polyimide thin-film electrode array was fabricated as a less invasive alternative to bone screws that could support a higher channel count. Electrodes, traces, and fiducials corresponding to bregma and lambda were all aerosol jet printed onto a polyimide substrate. Instead of drilling holes as was necessary for the bone screw ECoG, the array was implanted on the surface of the skull. A drop of 1x phosphate-buffered saline (PBS) was placed on the skull with a plastic pipette. The array was then placed on top of the droplet and aligned such that the bregma and lambda fiducials printed on the array coincided with (AP: 0.00; ML: 0.00) and (AP: -8.80; ML: 0.00) respectively. Once aligned, the PBS was removed with a swab, lowering the film to the skull surface.

To secure the film, a thin layer of clear C&B Metabond was applied over the cranial window. The procedure then continued to intracranial DiSc array implantation with the slight modification of needing to drill through both the film and the cement instead of just skull.

### Surface Electrode Implantation Method C: Microwire ECoG

This was the tertiary surface recording method intended to reduce the time of surgery but is not recommended given the less stable ECoG recording. Only Subject 5 received this hardware. Prior to surgery, lengths of insulated 30-gauge silver plated copper wire were cut from a spool, partially stripped of insulation to expose 5/32” of wire, and then fitted with a depth stop 5/32” from the tip of the wire on the exposed side. An additional stainless-steel, size 0-80 cup point set screw was soldered to a separate piece of 30-gauge silver plated copper wire to serve as a reference lead. The opposite end of the reference lead’s wire was soldered to a metal pin.

During surgery, markings for 15 microwires were made to denote the locations of the microwires to be implanted. The exact locations have been provided as supplemental text in the appendix (Supplemental Item 1). Holes were drilled with a 0.9mm dental burr at each location. An additional hole was drilled with a 1.35mm drill bit to house the bone screw reference lead. The bone screw was then driven into its hole with a hex driver. Each microwire lead was then placed in its respective hole and temporarily secured with two-part silicone (Kwik-Sil).

### Intracranial DiSc Array Implantation

Two additional holes were drilled in the skull at (AP: -2.76; ML: -1.80) and (AP: -6.12; ML: -4.00) with a 3/64” drill bit (Figure 1-B). A length of 0.042” OD polyimide tubing was cut and attached to the stainless-steel rod on the probe holder with an alligator clamp. The exact length of tubing varied between subjects so the target depth achieved is outlined in (Table 1). The tubes were moved to the holes using the arms of the stereotaxic frame and lowered to the surface of the skull. Petroleum jelly was used to insulate any gaps between the tubing and hole. Once sealed, both the polyimide tubes and the ECoG / EEG leads were secured to the skull using opaque C&B Metabond dental cement. Once dry, the alligator clamps were removed and the probe holder was lifted, leaving the polyimide tubing attached to the skull with an open lumen.

In subjects with ECoG surface electrodes, a 500um tungsten carbide guide wire with a polyimide depth stop 19.5mm from the distal tip was then passed into each of the polyimide tubes to create a pilot hole down to the implantation depth (Table 1), followed by a 700um PTFE-coated stainless-steel guide wire with a depth stop in the same location (n=4). In subjects that received a sub-scalp EEG grid, a single 800um guide wire was used to make the pilot hole (n=2) and generate a tapered tract for the device. Two 64-channel DiSc arrays with depth stops 19.5mm from the distal tip were then implanted by hand, one in each of the implanted polyimide guide tubes. The devices flanked the hippocampus from both sides on the anterior-posterior axis (Figure 1-D). Two-part silicone was used to secure the depth stops to the guide tubes and prevent radial motion. A picture was then taken from above so that the rotation of the PCB devices could be calculated during contact reconstruction.

### In Vivo Electrophysiology

DiSc microelectrode arrays were wired directly into an Intan RHD2000 recording system using a separate SPI cable for each device. For ECoG, pins soldered to the terminal ends of wires connected to each lead were routed into a 16-channel head-stage via a third SPI cable connected to an 18-pin electrode adapter board. For EEG, a 32-channel adapter was used to connect the film to a 32-channel head-stage. Recordings were taken using the Intan RHX Acquisition Software. Impedance measurements were taken for all implanted electrodes prior to beginning recordings using a 1kHz test frequency. The isoflurane percentage was set to 2% and then a recording was taken of the subject’s baseline neural activity for 30 minutes.

Once complete, the syringe containing KA was lowered to an adjusted injection depth of (DV: -9.93mm) while still at a 30-degree pitch. A new recording session was started and 300nL of KA was injected into the BLA at a rate of 100nL/min using an electronic injection syringe. If spike activity was not observed after 30 minutes of recording, another 300nL of KA was injected. After 30 minutes, if no synchronous sharp-wave events could be observed at a rate of 3Hz or faster, another 300nL was injected. No subject received more than 900nL of KA and the total volume of KA administered is outlined in (Table 1). Recordings started simultaneously with the first KA injection.

### Tissue Processing

Upon completion of recordings, subjects were sacrificed with 5% isoflurane and cervical dislocation. The two-part silicone used to secure the DiSc arrays was cut away and the microelectrode arrays were removed. In Subject 1, sealed 794um OD polyimide fiducials with depth stops 19.5mm from the distal tip were then placed into the guide tubes to serve as fiducial markers for the implanted arrays. This was not repeated in any of the other subjects. Once subjects were no longer responsive, they underwent transcardial perfusion with 200mL of cold PBS followed by 200mL of cold 4% paraformaldehyde in PBS. Subjects were decapitated via guillotine, and the entire head was soaked in 4% paraformaldehyde for 48 hours. Following the overnight soak in 4% PFA, skin and muscle tissue were sheared from the subject’s head, leaving the exposed skull with the brain inside. The cement was then removed from the skull using a hand rotary.

### Magnetic Resonance Imaging

Post-mortem T2 3D MRI images were taken using a 7T Bruker Bio-spin MRI. Perfused brain tissue was imaged in air with the subject’s muscle tissue removed and skull intact. Images were taken at a resolution of 200x400x200 voxels with an intervoxel spacing of 100um x 100um x 100um. Images were obtained using a 3D scan configuration with a gradient echo pulse sequence (Figure 3-C).

**Figure 2.**
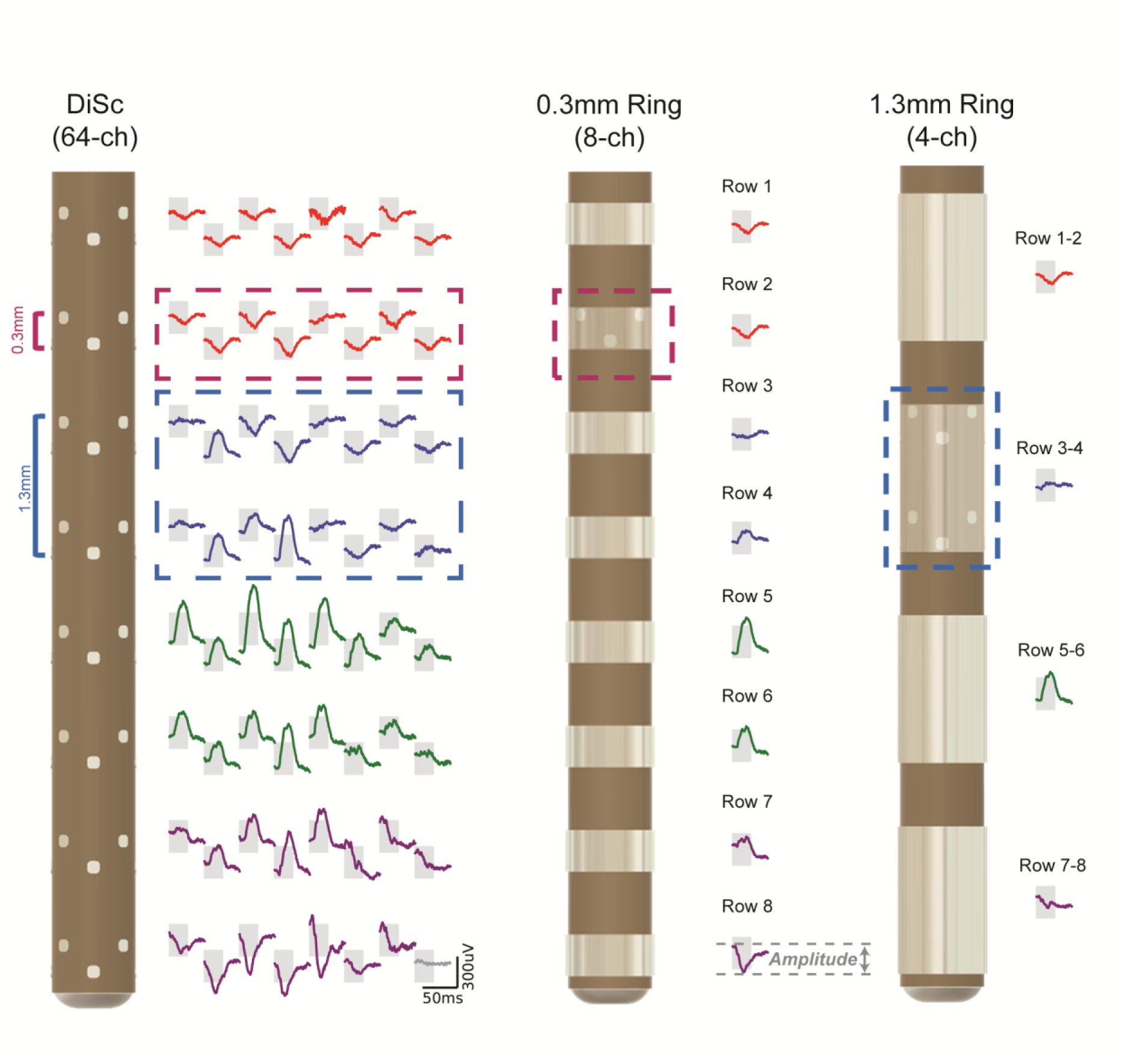
Virtual Signal Creation. Example signals from a DiSc microelectrode array (left), a 0.3mm ring electrode (center) and a 1.3mm ring electrode (right) during the same global time window. The 64-ch DiSc waveforms are drawn in a flattened rectangle as if the device were unwrapped. The grey box is a synchronized 25ms window across all channels. The bottom right trace (grey) represents the device blind/diagnostic channel and should be ignored. The DiSc signals were recorded in vivo, the virtual 0.3mm ring signals are the average of eight DiSc channels spanning one row, and the 1.3mm ring signals average 16 DiSc channels spanning two rows. The data shown are taken from Subject 1’s post-KA recording. Comparison of peak amplitude across electrode types was performed by measuring the maximum amplitude (peak-to-peak) for a distinct spiking event for each electrode type.

**Figure 3.**
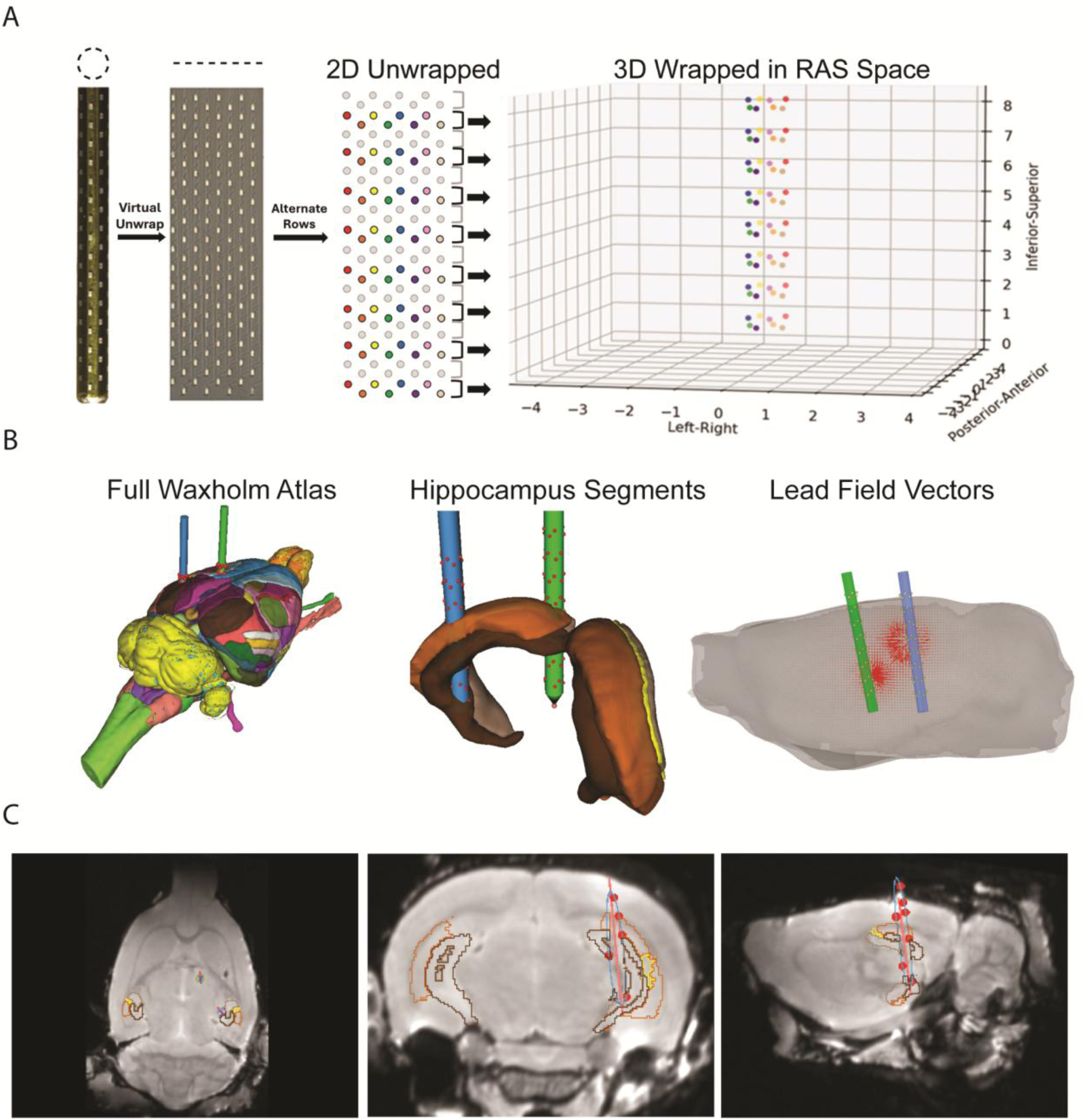
Virtual Reconstruction of Acute Surgery. (**A**) Virtual models of the implanted probes integrate design schematics and high-resolution micrographs of the fabricated devices to determine each channel’s location relative to the PCB. Electrodes can be plotted in a 2D plane or 3D coordinate space. Only 64 of the 128 channels on each device had an amplifier connection; only the active electrodes are shown. (**B**) Reconstructions of two DiSc devices in the full Waxholm Space Atlas (left), and hippocampal segmentations—dentate gyrus, CA3, CA2, CA1 (center). The atlas was normalized to the Subject 3 MRI and includes device tracts. Lead field vectors generated in Brainstorm for use in the inverse model of Subject 3 (right). (**C**) Cross-sections of T2 MRI from Subject 3 with hippocampal atlas segmentations as outlines. Red circles denote microelectrodes that are in-plane. Blue outline denotes cross- section of the posterior DiSc device that is in plane. Axial (left), coronal (center), and sagittal (right) views are shown.

### Data Processing

Data were processed offline using a combination of Python, MATLAB, Brainstorm, ImageJ and 3D Slicer. The raw recordings from the DiSc arrays were taken at 20kSamples/sec.

### Electrode Screening

DiSc configuration voltage signals were screened for both impedance and noise. Impedance measurements taken at the time of recording were used to exclude channels with an impedance over 1.5 MΩ. Ten seconds of data from the pre-injection baseline recording were then analyzed and any channels with a root mean square value that exceeded the data’s mean plus 2 standard deviations were omitted. Channels that satisfied both criteria were used in the subsequent steps.

### Virtual Ring Electrode Signal Creation

Voltage signals from DiSc channels that passed impedance and noise screening were spatially averaged to create virtual ring electrode signals (Figure 2; Supplemental Figure 2). Averaging was previously demonstrated to provide an amplitude within 5% of a full ring contact (Abrego *et al*., 2023). For these experiments, the DiSc electrodes were arranged in 8 diamond shaped rows around the device and every other ring of 8 was skipped. The first virtual ring configuration, the 0.3mm ring electrode, was created by averaging together all good channels in a row of 8 diamond-patterned electrodes. A second virtual ring configuration, the 1.3mm ring electrode, was created by combining two rows of 8 diamond-patterned electrodes – 16 DiSc electrodes in total. The ring electrodes typically used in clinical settings have a 2mm span; however, given the small size of Sprague Dawley rats in comparison to humans, the largest ring size used in this study is the 1.3mm ring.

### Data Preprocessing

After the virtual ring configuration signals were generated, all signals were preprocessed using the MATLAB implementation of Brainstorm. The data were bandpass filtered with a Kaiser FIR filter from 0.5 to 119Hz to isolate LFP activity while avoiding 120Hz as it is a harmonic of 60Hz power-line noise. Eighth-order IIR notch filters were then applied at 60Hz and 120Hz to further attenuate power-line noise. The data were then resampled from 20kHz to 2kHz to reduce computational complexity.

### Line Length Ratio

To further reduce the computational load, data from each device configuration were resampled to 1kHz for this analysis only. Line length was calculated according to (Equation 1) as outlined in other works (Esteller *et al*., 2001; Esteller, Echauz and Tcheng, 2004),

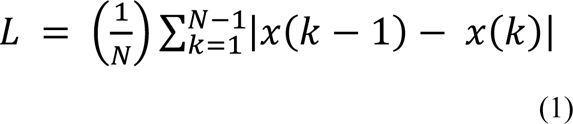

where L is the line length value, N is the number of samples in the window, and x(k) is the value of voltage signal x at sample k. For each signal, the absolute value of the differences between values were added over a one second discrete time window – a duration that has been shown to be effective in other works (Sun *et al*., 2022). This was repeated over the entirety of the baseline and post-KA injection recording sessions.

For each channel of each device configuration, the line length values calculated during the entirety of the baseline recording session were averaged together to generate one mean baseline line length value for the entire recording. The line length signals generated from both the baseline and post-KA injection recordings were then divided by the mean baseline line length to form a line length ratio (LLR) as defined by (Equation 2).

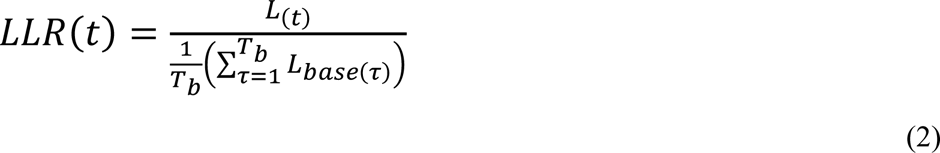

Here, LLR is the line length ratio, t is time in the current recording, L is the line length of the current window (Eq. 1), L_base_ is the line length of samples in the baseline recording, τ is time in the baseline recording, and T_b_ is the duration of the baseline recording.

### Automated Detection

To evaluate the spatial sensitivity of each device configuration, the LLR was used to determine when brain activity exceeded normal limits. An LLR greater than 1.75 has been reported as a commonly used threshold for clinical detection of seizures (Sun *et al*., 2022). To avoid false detections caused by noise, a 30-second time threshold was added as a secondary criterion for seizure detection. Thus, instances where a channel’s LLR exceeded 1.75 for at least 30 seconds were treated as successful detections.

Since the 1.3mm ring electrode has the largest contact size, it was used as the basis of defining zones to analyze in the tissue. The surface area of tissue in contact with each virtual 1.3mm ring also contacts two 0.3mm rings and 16 DiSc microelectrodes since the virtual signals were made via spatial averaging and the devices occupy the same volume (Figure 6-A). Each implanted array can fit 4 of these zones such that none of them overlap and no electrodes are counted twice. On each device, the post-injection LLR of the one 1.3mm ring, two 0.3mm rings and 16 DiSc electrodes that occupy the same area were evaluated against the previously described LLR and time thresholds.

**Figure 4.**
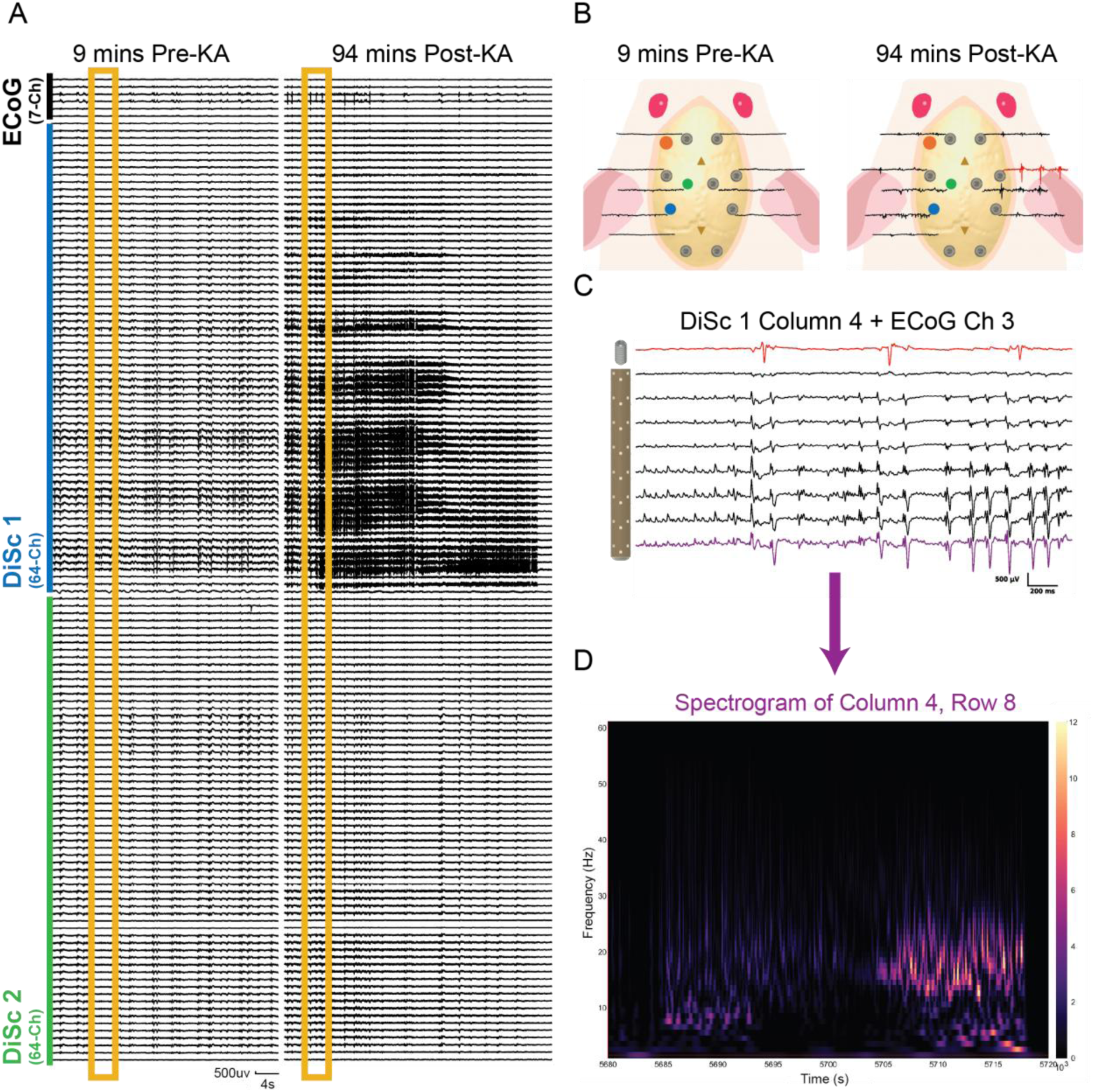
Effect of Kainate Injection. (**A**) Example of concurrent LFP recordings taken from two DiSc arrays and seven ECoG screws in Subject 1, nine minutes before (left) and 94 minutes after (right) the KA injection. Each plot represents a 40-second window of signals from all channels. The three-seconds of data shown on the skull plots in B are marked with an orange box. (**B**) Waveforms representing three seconds of data from the ECoG leads spatially arranged on a diagram of the rat skull. The top 2 rows of each DiSc electrode were also averaged together to simulate additional ECoG-like signals at the sites of implantation. Data are also from Subject 1 before (left) and after (right) the KA injection as denoted by the boxes on A. (**C**) Signals recorded from the ECoG lead above the contralateral dorsal hippocampus (top) and the eight channels in column four of DiSc 1 in the ipsilateral ventral hippocampus (bottom). All traces are from the three-second window denoted on the post-KA plot in A. (**D**) A spectrogram of the most ventral electrode of DiSc 1 column 4 over the same 40 second period shown in A.

**Figure 5.**
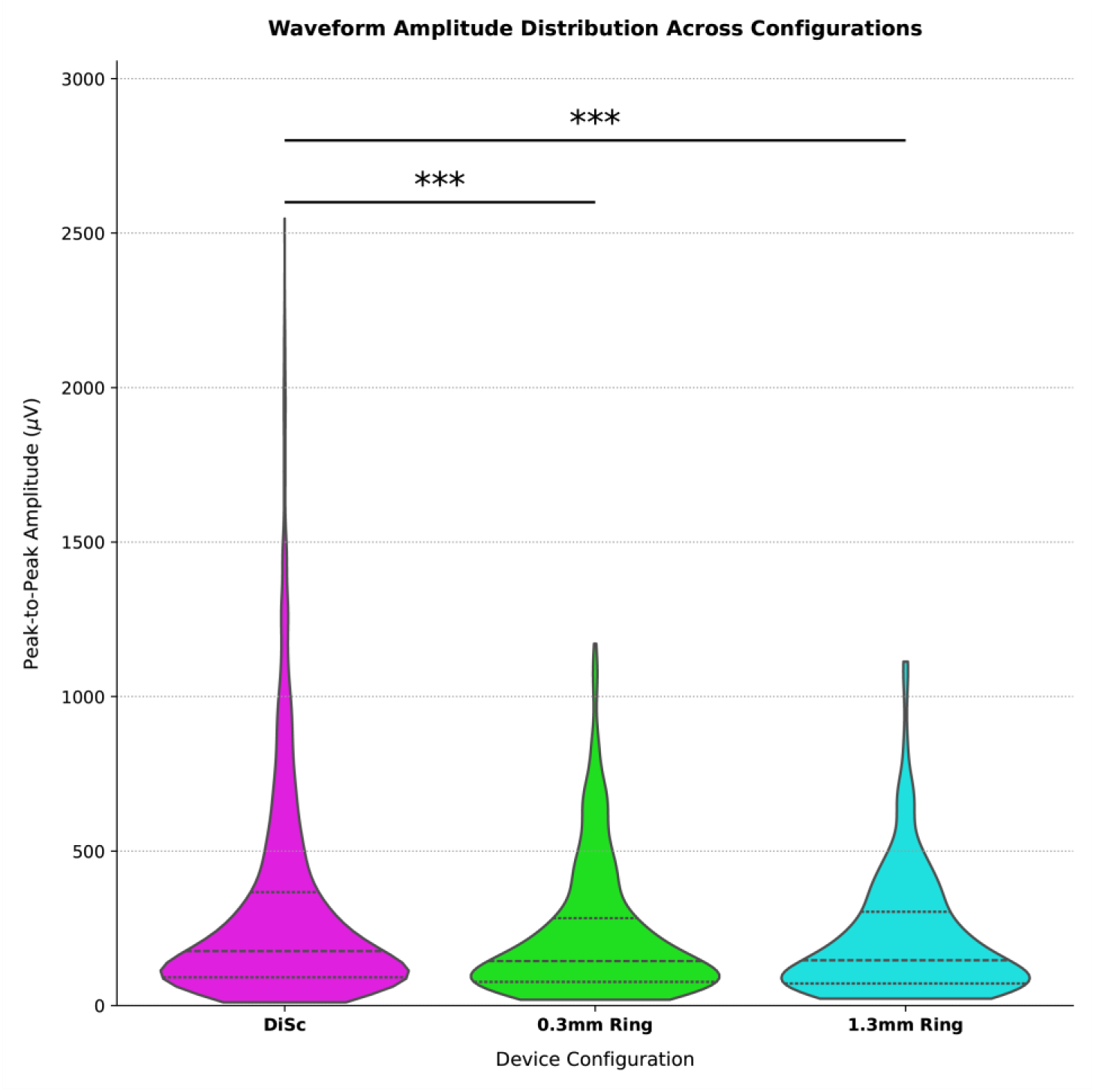
Peak-to-Peak Amplitude of Select Events. The peak-to-peak amplitudes of 10 spike events selected from periods of high LLR are compared between device configurations. A video showing the window from which Subject 1’s events were selected is available in the appendix (Supplemental Figure 7). DiSc configuration yielded significantly higher amplitudes than the 0.3mm ring by +82.6 µV (*β* = 82.639; 95% CI: 57.944-107.344; p<0.001, LMM) and 1.3mm ring by +93.9 µV (*β* = 93.884; 95% CI: 60.070-127.699; p<0.001, LMM) on average in subjects with KA seizures (n=4).

**Figure 6.**
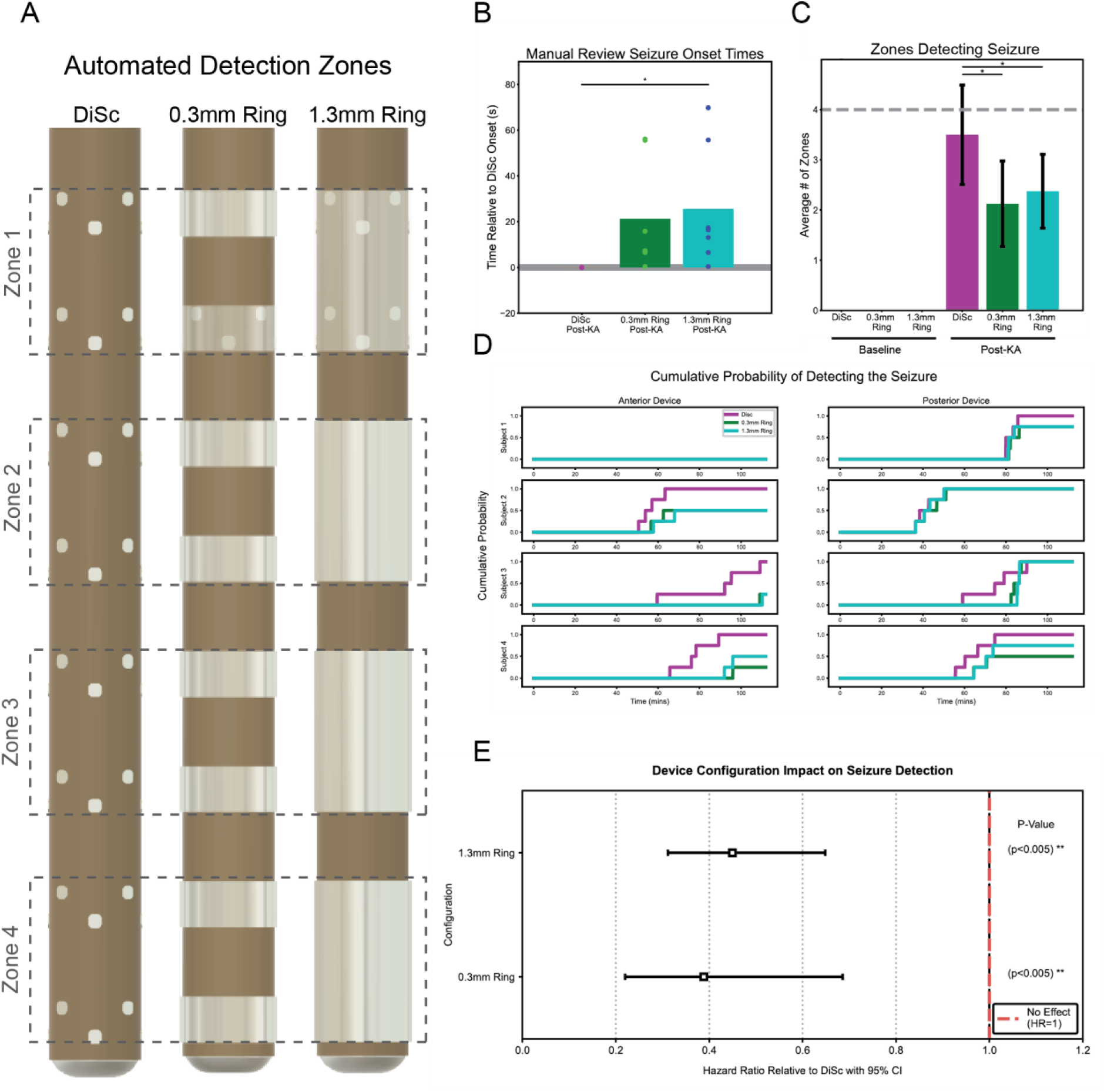
Manual and Automated Review of Seizure Data. **(A)** Example of 1.3mm zone in each configuration. These zones were only considered in the automated detection analyses and not the manual neurologist review. (B) Results of manual review of Analysis Group 1 from a board-certified epileptologist. Onset was first visible in DiSc configuration in all cases, so latencies for each configuration are expressed relative to the corresponding DiSc configuration onset time. All DiSc times are zero since they are the comparative time points. Bars show the mean value while individual onset latencies are overlaid as scatter points. The 1.3mm ring configuration had a significantly longer latency than the DiSc array (p=0.042). (**C**) Average number of 1.3mm zones per device that detected a seizure. The maximum possible value is four (grey line). No zones satisfied the detection criteria during their pre-injection baseline recordings so the value for each baseline group is zero. (**D**) Kaplan-Meier curve of the cumulative probability of a zone detecting a seizure. Steps in the plot represent a unique 1.3mm zone having its first seizure detection. All device configurations are plotted separately for each implanted device. (**E**) Scientific tree plot of the hazard ratios generated by the Cox Proportional Hazard analysis. The 95% confidence intervals are plotted as error bars. DiSc configuration was 2.56 times as like to have a new detection than a 0.3mm ring (HR = 0.39, 95% CI = 0.22-0.69, p<0.005) and 2.22 times as likely to have a new detection than the 1.3mm ring (HR = 0.45, 95% CI = 0.31-0.65, p<0.005)

To evaluate differences in the average number of zones that detected a seizure per device (bounded count integers from 0 to 4), data were modeled with a Generalized Estimating Equation (GEE) using a binomial family framework and a logit link function (Liang and Zeger, 1986; Zeger and Liang, 1986). Since each subject received a cluster of two physical implants, observations within the same brain cannot be treated as statistically independent. To prevent pseudoreplication, an exchangeable working correlation structure was enforced at the individual animal level, and standard errors were calculated using the robust Huber-White sandwich variance estimator (Hardin and Hilbe, 2012).

The total number of zones that detected a seizure by the end of each recording were calculated for each device. The zone counts were expressed in two columns – one with the number of zones that did detect the seizure, and one with the numbers of zones that did not. The sum of the two values in any row was always 4. The data were then grouped by subject ID and fit to the GEE model. The average number of zones that met the detection criteria were then plotted with error bars denoting the standard error (Figure 6-C). The odds ratios (OR) generated from the resulting coefficients are also reported.

### Time-to-Event Analyses

Time-to-event latencies were calculated to determine how long each zone took to have at least one channel meet the detection criteria after the start of the KA injection. Latencies were calculated at a time resolution of 1 second. A survival function, S(f), can be derived from these latencies by using the Kaplan-Meier method to estimate the unadjusted probability of a zone “surviving” – or in this case failing to detect a seizure – at each time point (Clark et al., 2003; Hosmer, Lemeshow and May, 2008; Schober and Vetter, 2018). Since the probability remains constant between events, S(f) is a step function where a vertical drop signifies the occurrence of an event (Bewick, Cheek and Ball, 2004). These functions can be plotted upwards or downwards to display the same information (Pocock, Clayton and Altman, 2002; Dudley, Wickham and COOMBS, 2016). To make the graphs more intuitive, the original downward survival curve was converted to an upward curve that shows the cumulative probability of having a zone detect a seizure (Figure 6-D). This was achieved by plotting the inverse of the survival function, 1-S(f), at every point to flip the graph vertically. Separate curves were plotted for each implanted array in all 3 device configurations.

To assess the effect of device configuration on seizure detection, a Cox proportional hazard model was fit to the time-to-event latencies. This method was chosen since some zones never reached criteria and thus introduced right-censoring to the dataset that would potentially bias other statistical methods. Observations were clustered by subject to adjust the standard errors and account for the fact that some measurements were nested within the same animals. The exponentiated coefficient, also referred to as the hazard ratio, was extracted and plotted on a scientific forest plot with the lower and upper 95% confidence intervals added as error bars (Figure 6-E).

### Amplitude Analysis

To compare the effect of device configuration on signal amplitude, 10 events were selected from a period of hyperactivity in each subject and the peak-to-peak amplitude recorded by each channel was calculated. The time window from which these events were extracted in Subject 1 is shown in a video in the appendix as an example (Supplemental Figure 7). These values were then used as input to a linear mixed effects model. Linear mixed effects models are an effective means of analyzing data that are clustered or derived from repeated measures (West, Welch and Galecki, 2022). Bad channels were excluded as previously described.

The amplitude in microvolts recorded by each good channel at each event time was used as the model’s dependent variable. The fixed effect to be tested for significance was the device configuration. Due to the nested structure of the dataset, the subject ID and a categorical label for implantation position – anterior or posterior – were treated as random effects to cluster amplitudes based on the conditions under which they were recorded. This information was processed with a linear mixed model using the DiSc configuration data as the reference set. The results of the model fitting were an intercept denoting the mean amplitude and categorical coefficients denoting the expected relative signal change in microvolts that results from switching to each of the ring configurations. Violin plots of the peak-to-peak amplitudes were then generated (Figure 5), and the resulting coefficients are reported.

### Virtual Reconstruction

The positions of each device configuration’s contacts were reconstructed in 2D and 3D space to aid in analysis of the spatial character of the data obtained.

### Contact Reconstruction

The manufacturing CAD used to fabricate each DiSc device was measured in the software KLayout to measure contact locations in 2D space. Prior to implantation, each DiSc device was also measured under a microscope to obtain the shaft diameter, rate of helical twist, and angle of the electrode array’s seam relative to the printed circuit board (PCB) back end. DiSc is effectively a flat microelectrode array wrapped around a cylindrical core so signals can be arranged in 2D space based on specifications of the unwrapped array. The coordinates of each contact in the device schematic were arranged in a plane (Figure 3-A). The profile of the DiSc array was assumed to be a perfect circle, so the horizontal coordinate of each electrode was divided by the circumference of the real probe to convert the horizontal positions to percentages. These percentages were converted to radians by multiplying them by 2π. The helical twist and PCB and angle were then added as offsets in radians.

The final rotations were converted from polar coordinates back to 2-dimensional cartesian coordinates in a flat circle, creating the x-axis (left-right) and y-axis (anterior-posterior) coordinates for the 3D reconstruction. The vertical positions of each contact on the flattened 2D device schematic were then used as the initial z-axis (inferior-superior) coordinates. An offset accounting for the distance from the tip of the DiSc probe core to the lowest electrode was then used to translate the coordinates such that the device tip was the origin of the coordinate space (Figure 3-A). Similar to how voltage signals from the DiSc arrays were spatially averaged to create virtual ring electrode signals, the locations of the virtual rings were calculated by averaging the 3D coordinates of the DiSc electrodes that contributed to them.

### Line Length Ratio Heat Maps

Heat maps were generated by combining the 2D electrodes’ coordinates with the calculated LLR values (Figure 8). Each heatmap shows the mean LLR value of each channel over a 5-minute window. This longer time window was chosen to find regions that were stably and consistently active. The values were spatially arranged using the 2D coordinates measured from the design specifications (Figure 3-A). For DiSc heatmaps, inverse distance weighted interpolation was applied to the values to account for the areas between the sampled electrode locations. The resulting values were then grouped into 20-level contours to articulate their shape. Using the 2D to 3D conversion, the 2D heatmaps could also be represented in 3D cylindrical form (Figure 3-A). For ring electrode heatmaps, linear interpolation was used as the contacts extend along a single axis (Figure 8-B,C).

**Figure 7.**
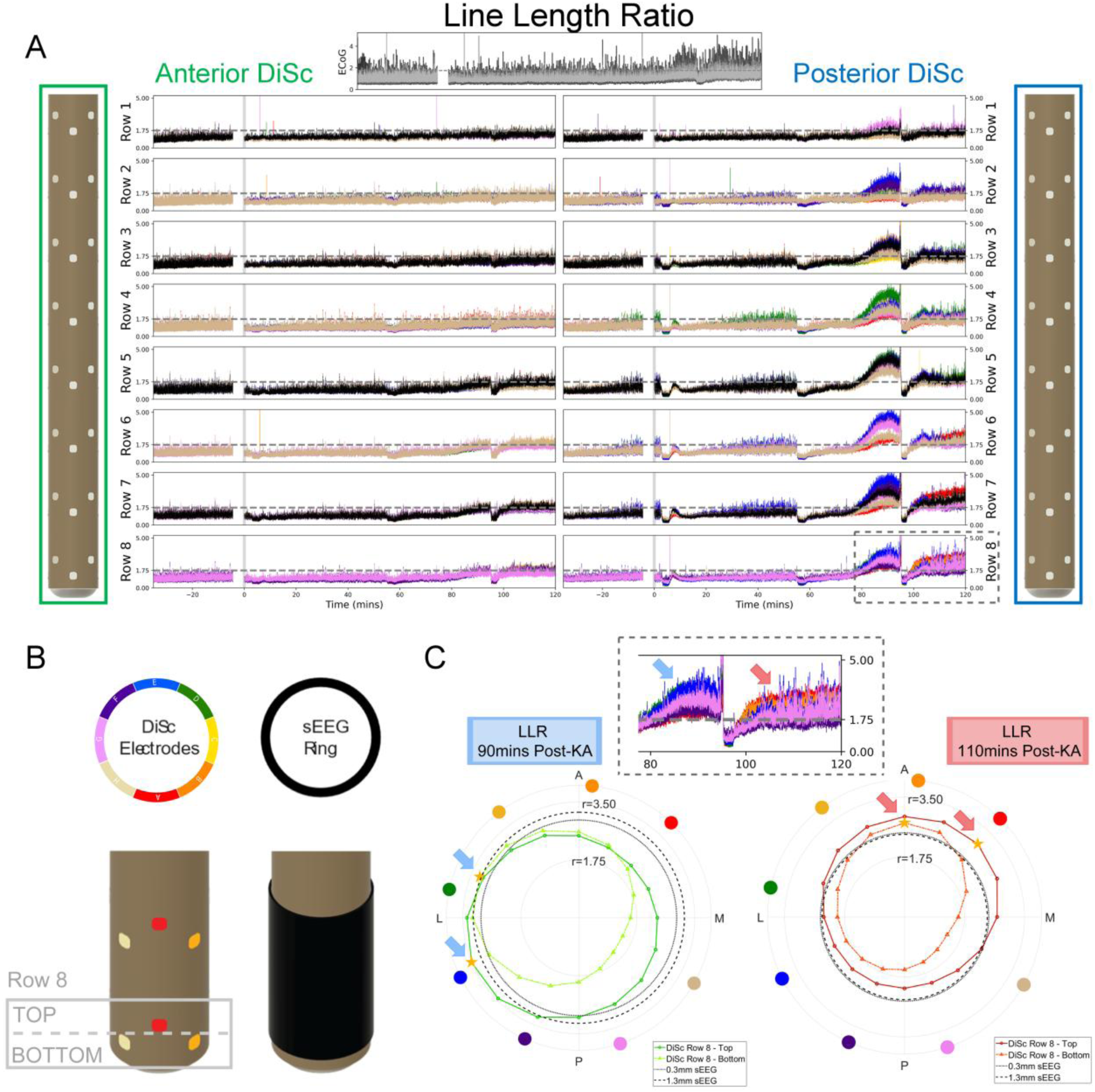
Line Length Ratio (LLR) (**A**) LLR computed for Subject 1. Each virtual 1.3mm LLR signal was created from DiSc electrodes from two rows and overlaid on the top row (black, every other row). The dotted box on posterior DiSc row eight denotes where the data shown in Figure 7-C was sourced. A horizontal dotted line spans each graph to represent the 1.75 LLR threshold. A vertical solid line denotes the beginning of the kainate injection. (**B**) Signal direction shown by rainbow coloring convention as you move around the device. The 1.3mm sEEG rings are black. (**C**) Polar plots of the LLR at 90 (left, blue) and 110 mins (right, red) post-KA injection to illustrate source movement as seizures evolved. The top and bottom 4 contacts from row 8 are 230 microns apart and illustrate orientation differences across a small gap. Maximum LLR value marked (star and arrow).

**Figure 8.**
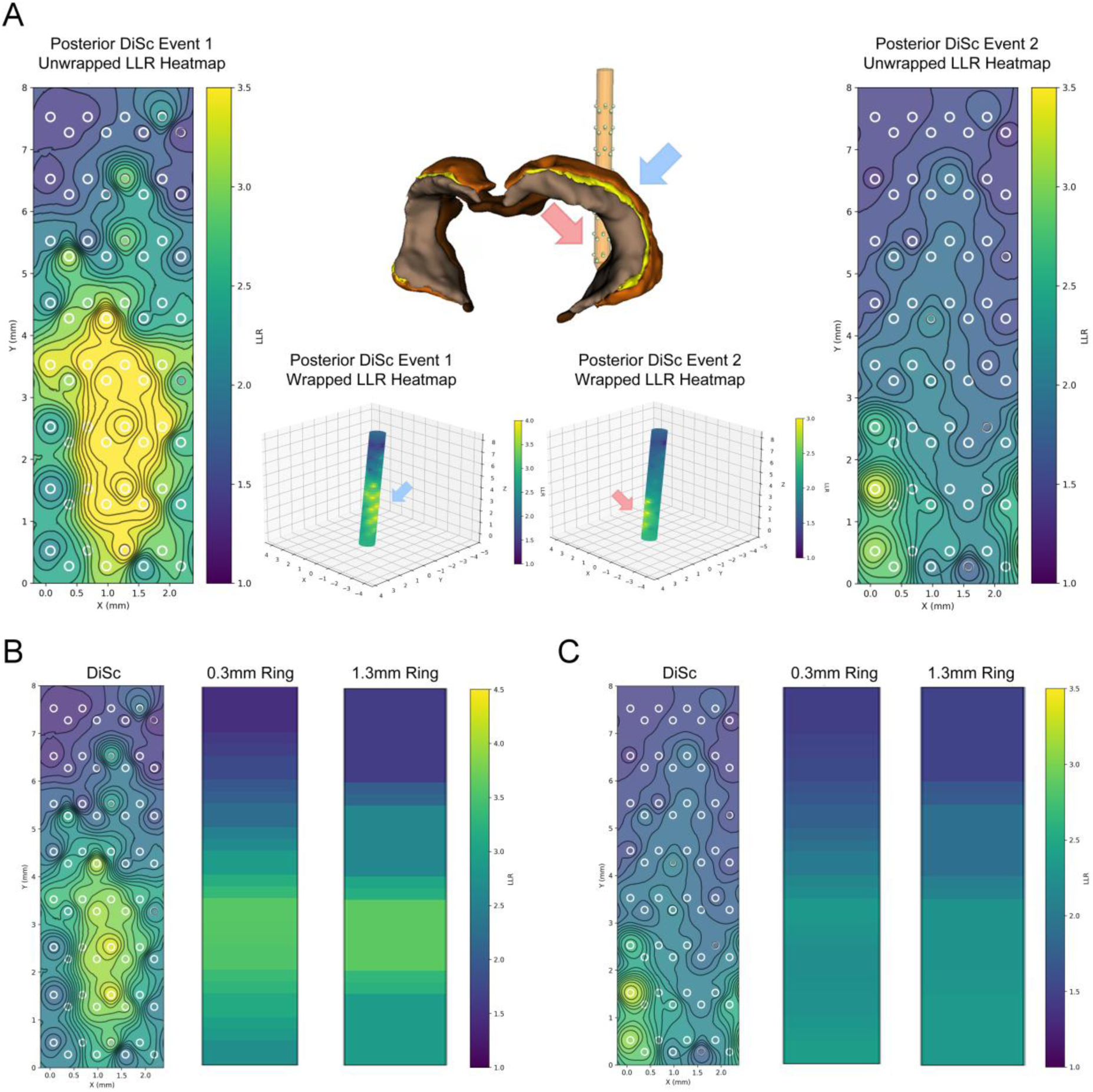
Line Length Ratio Heatmap. (**A**) Heat maps showing the mean LLR over 5 minutes at two different time points (far left, far right) in the post-KA injection recording from Subject 1. The x- and y-axes denote position on the flattened array in millimeter coordinate space. 3D heat maps of the same events are also shown (center bottom). Virtual reconstruction of the posterior DiSc device is shown in the Waxholm atlas hippocampal segmentations (dentate gyrus, CA3, CA2, CA1) is registered to the MRI of Subject 1 (center top). (**B, C**) Heatmap representation of events 1 and 2, respectively, using a common heatmap across device configurations. The heatmap scale has been extended in B to better show the contour.

### Trajectory Alignment

The skull and surrounding tissue were digitally removed from each subject’s MRI in Python before placing them into a 3D coordinate space created in the program 3D Slicer. A mask of the brain was created from the MRI and exported as a 3D model for use in the inverse modelling. A copy of the Waxholm Space Atlas of the Sprague Dawley Rat Brain (Papp *et al*., 2014; Kleven *et al*., 2023) was normalized to the subject’s MRI in Python and imported to the 3D Slicer environment. The virtual electrode positions were placed into the same coordinate space as their respective subject’s MRI (Figure 3-B).

Alignment of the virtual devices to the subject MRI was accomplished by writing a custom 3D Slicer extension. The tracks of the implanted DiSc devices were still visible in the MRI after the devices were removed. Two points along the visible tract were selected and used to generate a line representing the device’s trajectory. The coordinate space of the 3D electrode positions was aligned to the trajectory using Rodrigues’ rotation formula with an additional translation. A separate rotation parameter was added to allow the device to be twisted along its trajectory (Figure 3-C). ImageJ was used to calculate the rotation of the device relative to the horizontal axis of the stereotaxic frame using photos acquired during the acute surgery. The angle was then input to the twist parameter and used to rotate the virtual devices. The final electrode locations were then exported and transferred to Brainstorm.

### Brain Source Localization

Source localization was performed in Brainstorm (Tadel et al., 2011) using a minimum-norm estimation (MNE) framework with sLORETA normalization (Baillet, Mosher and Leahy, 2001; Pascual-Marqui, 2002). Forward and inverse solutions were computed using subject-specific MRI anatomy and electrophysiological recordings within the Brainstorm-DUNEuro finite-element modeling pipeline for realistic head modeling and source estimation (Medani et al., 2023). The resulting framework enabled computation of lead fields for the DiSc electrode geometry and subsequent inverse modeling of seizure-related field activity.

The reconstructed electrode coordinates, registered MRI volumes and Waxholm atlas segmentations described in the previous section were subsequently imported into Brainstorm (Tadel et al., 2011) to generate subject-specific anatomical models for forward and inverse source estimation. Masks of the brain volume were also generated in 3D Slicer and exported as triangulated surface representations for volumetric meshing. Within Brainstorm, geometrical representations of the DiSc electrode shafts were generated and embedded into the volumetric brain model prior to tetrahedral mesh generation. The extracted brain surfaces were smoothed to reduce discretization artifacts and converted into tetrahedral volumetric meshes using Iso2Mesh for finite-element modeling (FEM). Mesh density was locally refined in the vicinity of implanted DiSc contacts to improve numerical stability and spatial resolution of the forward solution.

The final FEM model consisted of a smoothed brain volume modeled as a homogeneous conductive medium together with the two implanted DiSc depth electrodes and their associated insulating shaft geometries. Brain tissue was assigned a homogenous conductivity value of 0.33S/m while the DiSc electrodes shafts were modeled using a low conductive value of 1e-5 S/m. A volumetric source space was defined throughout the brain using a regular 0.2 mm isotropic grid.

Forward modeling was performed using the FEM implementation provided by DUNEuro, integrated within Brainstorm (Medani *et al*., 2023, 2025). Lead-field matrices were then computed separately for DiSc, 0.3mm ring and 1.3mm ring configurations. To improve numerical stability of the FEM computation and avoid spurious lead field values, an exclusion zone of 0.5mm was applied around each contact such that source points located within 0.5 mm of the contacts were excluded from the source space. Preprocessed voltage time-series were imported into Brainstorm and aligned with the corresponding electrode geometries for each device configuration. Noise covariance matrices were estimated from baseline recordings acquired prior to KA administration. Covariance regularization was performed using Brainstorm’s median eigenvalue regularization approach.

Distributed source estimates were computed using a minimum-norm inverse solution with sLORETA normalization applied exclusively to sEEG depth channels; surface ECoG channels were excluded from the inverse model. Source reconstructions were computed independently for each device configuration to enable direct comparison of spatial dispersion and directional specificity across electrode geometries. The resulting volumetric reconstructions were visualized in Brainstorm and exported for cross-configuration comparison and analysis (Figure 9; Supplemental Figures 4–7).

**Figure 9.**
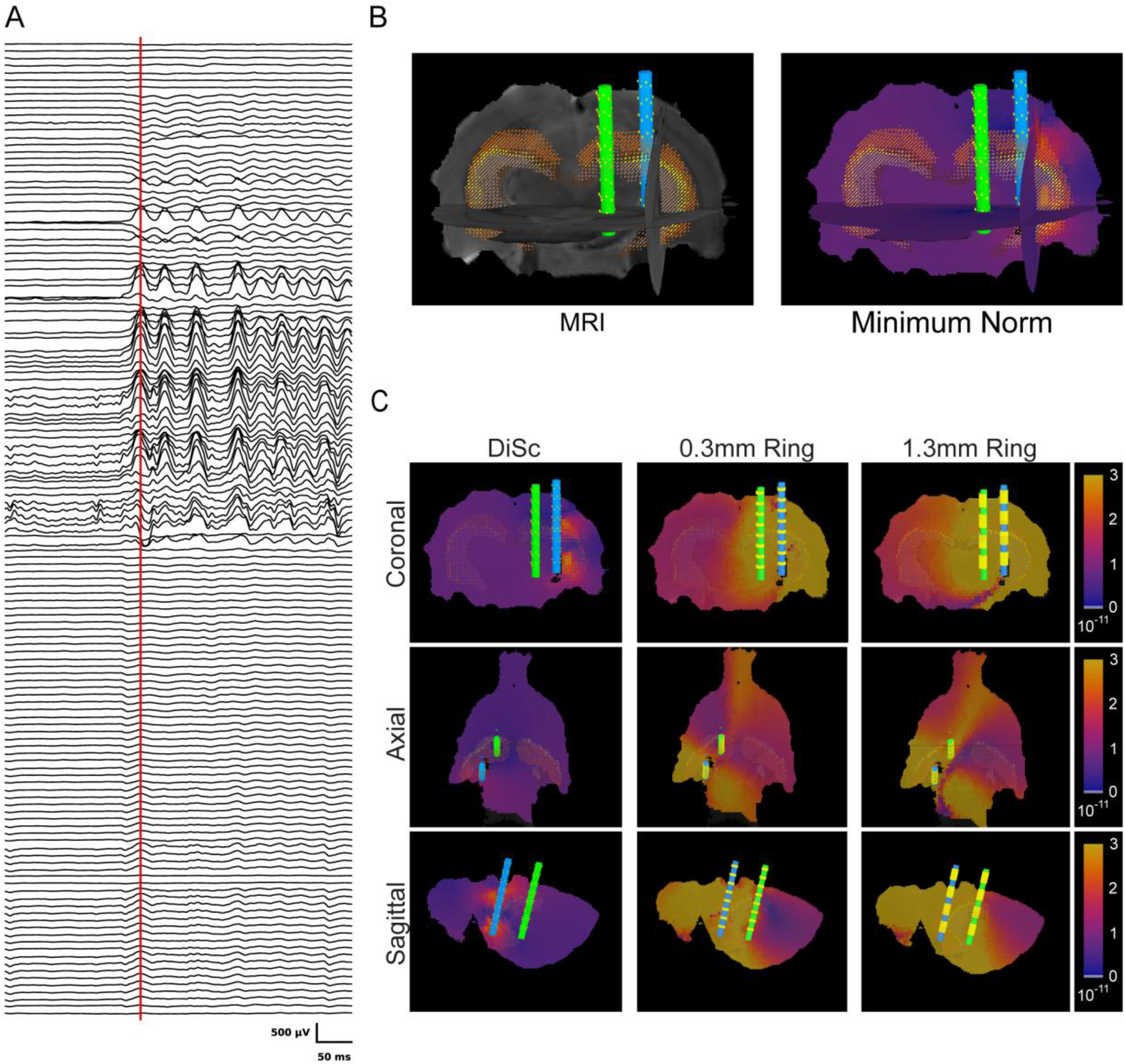
Minimum Norm Projection. (**A**) Signals recorded from the DiSc arrays. The red line denotes the sample used to generate figures B and C. (**B**) The MRI of Subject 1 with normalized Waxholm Atlas segmentations overlaid (left) and the sLORETA minimum norm projection 5699.695s after the kainate injection (right). Hippocampal subregions – dentate gyrus (dark brown), CA3 (light brown), CA2 (yellow), CA1 (orange) – are shaded in. (**C**) The sLORETA minimum norm projection at 5699.695s post-injection in Subject 1. A slice of the result for each device configuration is shown along the coronal (top), axial (center), and sagittal (bottom) axes.

## Results

### Acute KA Injection

Simultaneous extracranial and intracranial EEG recordings were taken under isoflurane anesthesia in Sprague Dawley rats (n=6) both before (mean duration: 31.20 min, range: 30.05-36.00 min) and after (mean duration: 121.47 min, range: 111.08-130.62 min) an intracranial KA injection in the BLA (Figure 4-A,B). At the time of surgery, sharp events with a spatially coherent falloff were observable in signals recorded from the microelectrode arrays and surface electrodes (Figure 4-C). A power increase that evolved over time was also observed in LFP bands (Figure 4-D).

### Signal Amplitude

Relative to the virtual ring electrodes, DiSc configuration yielded a significant improvement in signal amplitude. Linear mixed-effects modeling confirmed an average increase in peak-to-peak amplitude of +82.6 µV (*β* = 82.639; 95% CI: 57.944-107.344; p<0.001) compared to the 0.3mm rings, and +93.9 µV (*β* = 93.884; 95% CI: 60.070-127.699; p<0.001) compared to the 1.3mm ring (Figure 5). To determine if the increase in spiking observed in the data was driven by noise, a noise floor was calculated using a pre-surgical recording in PBS. In the subjects that showed signs of seizure activity post-injection, Analysis Groups 1 and 2 (n=4), presurgical noise recordings showed a slight trend of noise decreasing as electrode size increased but there was not a significant difference in noise between populations (Supplemental Figure 3).

### Epileptologist Review of Seizure Activity

To verify the presence of electrographic seizures, raw signals from the DiSc microelectrodes and virtual signals from 0.3mm and 1.3mm ring electrodes were preprocessed then reviewed by a board-certified epileptologist, JG. Inspection of the waveforms recorded from all subjects (n=6) identified a subset of rats (n=3) that showed seizure activity post-injection with a paroxysmal shift from baseline which make up Analysis Group 1 (Table 1). In a fourth subject, seizure activity was observed post-injection but without a paroxysmal shift from baseline due to the presence of spiking activity in the baseline recording (n=1) so it is classified as Analysis Group 2 (Table 1). In the remaining rats, notable seizure progression was not observed post-injection (n=2), creating Analysis Group 3.

In Analysis Group 1, JG manually identified seizure onset events visible in all device configurations (n=7). The latency of each onset relative to the start of the KA injection was calculated for each device configuration. Manual review of virtual signals from the 0.3mm ring electrodes yielded onset markings that were on average 21.22s later than when reviewing DiSc data (t(6) = 2.57, p=0.058, paired t-test). The 1.3mm ring electrode signals led to onsets being reported on average 25.60s later than the DiSc result (t(6) = 2.33, p=0.042, paired t-test). Reviewing data in DiSc configuration led to the earliest detection time in all cases (Figure 6-B).

### Automated Review of Seizure Activity

Subjects that showed seizure activity following KA administration, Analysis Groups 1 and 2 (n=4), also had LLR calculated for each device configuration. The latency to satisfy the automated detection criteria was evaluated in 4 non-overlapping 1.3mm long zones per device (Figure 6-A). Generalized Estimating Equation (GEE) modeling revealed a significant population-averaged effect of device architecture on seizure detection. Devices in DiSc configuration demonstrated the highest sensitivity to seizures with an average of 3.5 zones out of 4 that detected a seizure (an 87.5% success rate) during the post-injection recordings. Zones in DiSc configuration were 6.25 times as likely to experience an event (OR: 0.16; 95% CI: 0.03-0.86; p=0.033) than the 0.3mm ring configuration which had an adjusted mean of 2.125 zones out of 4 (a 53.13% success rate). DiSc configuration was also 4.76 times as likely to experience an event (OR: 0.21; 95% CI: 0.05-0.88; p = 0.033) than the 1.3mm ring which had an adjusted mean of 2.375 zones out of 4 (a 59.38% success rate). None of the devices had any electrodes satisfy the detection criteria during baseline recordings (Figure 6-C).

The time-to-event detection latencies of each implanted device were plotted as cumulative probability curves (Figure 6-D). The Cox proportional hazard was used to determine the relationship between the curves generated by each device configuration. The hazard ratios (HR) showed that DiSc configuration was significantly more likely to have a new zone detect the seizure event than both of the ring electrode configurations. The Cox proportional hazard showed that DiSc configuration was 2.56 times as like to have a new detection than a 0.3mm ring (HR = 0.39, 95% CI = 0.22-0.69, p<0.005) and 2.22 times as likely to have a new detection than the 1.3mm ring (HR = 0.45, 95% CI = 0.31-0.65, p<0.005) as is shown in (Figure 6-E).

### Virtual Reconstruction

The LLR signal over time is provided for Subject 1 (Figure 7) as well as Subjects 2-4 (Supplemental Figures 4-6). Virtual reconstruction allows for device signals to be laid out in 2- and 3-dimensional space. Subject 1 had two periods where the LLR exceeded 1.75 that were separated by a return to subthreshold activity (Figure 7-A). Using the increased spatial resolution of DiSc configuration to do 2D reconstruction revealed that the elevations in LLR had maximal values at locations that differed in both their depth and position around the circumference of the device (Figure 7-C). These locations mapped to the posterior device implanted in the ventral regions of the hippocampus (Figure 8-A). The 0.3mm and 1.3mm ring electrodes could only distinguish the differences in depth as they lack the radial sensitivity needed to evaluate direction (Figure 8-B,C).

### Minimum Norm Projection

Following virtual reconstruction, the minimum norm solution was evaluated for subjects in analysis groups 1 and 2 (n=4). Examples of the sLORETA result for each device configuration at the same time point are shown for Subject 1 (Figure 9) and subjects 2-4 (Supplemental Figures 4-6). A video showing the dynamics over time is also available in the appendix (Supplemental Figure 7). DiSc configuration yielded the most concentrated result volume. The 0.3mm and 1.3mm ring configurations yielded much more diffuse volumes with large values spread across the left hemisphere. The coordinates of the maximum confidence points in the DiSc result were then mapped back to a copy of the Waxholm Space Atlas that was normalized to an ex-vivo MRI of the subject’s brain. The points corresponded to the ventral hippocampus, specifically CA3 and the dentate gyrus.

## Discussion

Intracranial KA injections yielded epileptiform activity in Subjects 1 to 4. The presence of pre-injection spiking in Subjects 4 and 5 speaks to the importance of using devices with an appropriate form factor. While the 0.8mm diameter array is ideal for human surgery – the ultimate goal of this translational project – the small size of the rat brain increases the risk of damage during insertion. The rodent surgery must therefore balance between implanting the array deep enough to embed the entire 7.5mm span of electrodes but not so deep that injury potentials are generated. Subjects 4 and 6 were implanted deeper than the other subjects in an effort to ensure the topmost rows were embedded in cortex and not cerebrospinal fluid. However, both of these procedures saw abnormalities with Subject 4 having pre-injection spikes and Subject 6 not having epileptiform activity at all. Recent advancements in DiSc array manufacturing have enabled more flexible control over the selection of active electrodes. Should this study be repeated, an electrode layout that concentrates more channels near the distal tip of the probe should be used to reduce the necessary insertion depth.

The linear mixed model aimed to answer the high-level question of whether, across the events sampled, each of the ring electrode configurations capture smaller waveform amplitudes on average than the DiSc configuration. While the Cox proportional hazard was calculated with only a single level of clustering due to model limitations, the linear mixed-effects model can handle multiple clusters allowing implantation location to also be utilized as a random effect. Since peak-to-peak amplitude is a continuous real numeric measurement, a standard linear mixed model is sufficient. For discrete integer counts like the average zone calculation, a generalized linear mixed model or generalized estimating equation is more appropriate. The model reported that DiSc arrays provide a significant improvement to signal amplitude as was expected based on previous work using those devices (Abrego et al., 2023).

When manually reviewed by an epileptologist, data from the DiSc array configuration yielded the shortest onset latencies in all cases (Figure 6-B). The increased spatial resolution of the array results in the device sampling less tissue per electrode. In doing so, signal mixing can be greatly reduced which avoids the forced regression towards the mean that typically occurs in larger contacts. Such resolution can improve the clarity of low-voltage fast activity, a marker used in manual seizure detection, that would otherwise be neutralized by the other tissue in contact with the electrode.

However, while using microelectrode arrays improves the resolution of the tissue sampling, it also creates more data that needs to be reviewed. A standard ring electrode typically used for sEEG has 4-18 contacts spaced 2-10mm apart (Iida and Otsubo, 2017). Increasing the electrode density to 64 contacts per probe can create a data set that quickly becomes overwhelming as care providers often have to make rapid decisions while treating patients. Thus, effective means of dimensionality reduction that retain the spatial information but reduce the channel count back to quantities comparable to ring electrodes could prove incredibly useful in a clinical setting.

The DiSc electrode configuration also outperformed ring electrodes in automated detection using an LLR algorithm with thresholds. The disc arrays were able to sense an increase in LLR that satisfied the detection criteria in 87.5% of the areas sampled compared to the 0.3mm and 1.3mm rings at 53.1% and 59.4% respectively (Figure 6-C). Of the 8 devices implanted across the 4 subjects in Analysis Groups 1 and 2, DiSc reached threshold in all sampled areas for 7 of them. The ring electrodes only reached criteria in all sampled areas on two devices while the other five reached threshold in only a fraction of the areas. The Cox proportional hazard (Cox PH) model is the most commonly used survival data analysis technique for evaluating multiple covariates (Bradburn et al., 2003; Schober and Vetter, 2018). Cox PH doesn’t directly model the survival function but rather the hazard rate and thus can avoid the statistical biases caused by censoring. Since some of the devices did not detect the seizure in all 4 zones, the data set was right censored which necessitated its use.

DiSc configuration’s increased sensitivity opens the possibility of setting more strict detection criteria with higher thresholds, creating more accurate distinctions between seizure and non-seizure states. It is worth noting that the 1.3mm ring slightly outperformed the 0.3mm ring in the automated detection task – there were two areas in Subject 4 where the 1.3mm configuration reached criteria, but the 0.3mm configuration did not. However, DiSc configuration dramatically outperformed both ring configurations suggesting that increasing spatial resolution is more pertinent than trying to glean constructive effects from averaging signals from larger contacts.

One limitation to the LLR seizure detection analysis is the advantage potentially conferred by more channels for the DiSc device type. Decomposing a large sensor into smaller ones is our intent and many in the field have been hesitant to do because the smaller contacts produce greater a noise floor from the inherent Johnson-Nyquist noise, thus the scientific question should be “Which architecture gives the earliest or most frequent detection opportunities when all contacts are used?” It is not our intent to show that an equal number of micro contacts would have the same sensitivity as macro contacts.

Virtual reconstruction revealed the presence of high LLR events at similar depths with different spatial profiles. Due to their lack of radial sensitivity, conventional ring electrodes struggle to differentiate between the two as there is only a slight difference in LLR when signals around the entire circumference of the device are averaged together. In contrast, DiSc configuration provides both depth and radial information (Figure 8). Registration of the Waxholm Space Atlas to the subject’s MRI further improves the contextualization of the LLR signal as it is then possible to see what brain segmentations are in the vicinity. However, due to volume conduction in the brain, the atlas segmentation touching the electrode with the highest LLR is not necessarily the brain region generating the signal.

The minimum norm projection better accounts for this by utilizing inverse modelling methods including the conductive properties of the brain tissue. When applied to Subject 1, DiSc configuration’s minimum norm projection had the most concentrated sLORETA result distribution. The volume containing the highest confidence values increased as contact size increased with the 0.3mm ring configuration having more spread than the DiSc but less than the 1.3mm ring configuration (Figure 9). Achieving higher confidence in a smaller volume using directional depth arrays has potential to improve signal localization in patients undergoing epilepsy monitoring. We have not yet shown that high-density sEEG will be able to reduce the total number of implants per subject, but next steps include applying statistical and information theory to the localization problem. The use of DiSc arrays may prove to be especially useful when implanted at the edges of the hypothesized SOZ to identify borders with greater confidence or dodge critical vasculature while still contributing to localization efforts.

In conclusion, here we show that DiSc arrays can outperform ring electrodes in tasks relevant to seizure diagnosis. Microelectrode contacts achieved lower seizure detection latencies in both manual and automated detection. DiSc devices also detected increases in LLR across more of the area sampled by the probes. Large seizure events contained directional character that could be detected by microelectrodes wrapped on an insulating core. Ring electrodes are not sensitive to such differences, and their amplitudes represent regression to the mean. The minimum norm projection further emphasized the value of directional information as the DiSc configuration yielded a smaller spread in the sLORETA metric than the 0.3mm or 1.3mm ring configurations. Together, these results suggest that directional and scalable electrode arrays could improve seizure detection and localization efforts and contribute to the diagnosis and treatment of epilepsy.

## Data availability

All the data supporting this paper and the results will be available as a Brainstorm protocol within Zenodo (to be linked).

## Acknowledgements

Funding is from UTHealth Neurology and devices were developed and fabricated under work supported by NIH NINDS 1RF1NS133972 and 1UG3NS125487. We would like to thank the Rice University clean room staff. NT, CM, and JPS received funding from NIH NINDS UG3NS125487, NIH NINDS RF1NS133972, and Rice IITK global initiative funding. JCM received funding from NIH/NIBIB R01EB026299, NIH NINDS UG3NS125487, and NIH NINDS RF1NS133972. AAJ and RML received funding from NIH/NINDS R01NS121761, NIH/NIBIB R01EB026299, and DOD/ USAMRAA HT94252310149.

## Author Contributions

**Ryan Shores:** Conceptualization, writing – original draft, writing – review & editing, visualization, validation, supervision, software, methodology, formal analysis, data curation. **Takfarinas Medani:** Methodology, formal analysis, conceptualization. **Anand A. Joshi:** Methodology. **Camryn Matthews:** Methodology. **Yash Vakilna:** Methodology. **Jay Gavvala:** Review & editing, formal analysis. **Richard M. Leahy:** Resources, review & editing. **Sandipan Pati:** Conceptualization, resources, funding acquisition. **John C. Mosher:** Methodology, conceptualization. **Nitin Tandon:** Review & editing, resources, funding acquisition. **John P. Seymour:** Conceptualization, writing – review & editing, supervision, resources, methodology, investigation, funding acquisition.

## Supplementary Materials

### Supplemental “Figure” 1, Extra Methods Surgical Methods

During surgery, a burr hole was drilled at (AP: 2.37; ML: 4.9) with a 1.4mm dental burr to serve as the entry point of the KA syringe which would be lowered to (DV: -9.93). ECoG bone screw electrodes were implanted at (AP: 3.00; ML: 2.00), (AP: 3.00; ML: -2.00), (AP: - 2.76; ML: 1.80), (AP:-1.76; ML: 4.50), (AP: -1.76; ML: -4.50), (AP: -6.12; ML: 4.00), (AP: - 10.50; ML: -2.0) with an additional screw at (AP: -10.50; ML: 2.00) to serve as the reference electrode (Figure 1-D).

For subjects that received microwires, wires were implanted at (AP: 3.00; ML: -1.80), (AP: 3.00; ML: 1.80), (AP: 0.12; ML: -1.80), (AP: 0.12; ML: 1.80), (AP -2.76; ML: -4.00), (AP: - 2.76; ML: 1.80), (AP: -2.76; ML: 4.00), (AP: -6.12; ML: -1.80), (AP: -6.12; AP: 1.80), (AP: - 6.12; ML: 4.00), (AP: -8.31; ML: -4.00), (AP: -8.31; ML: -1.80), (AP: -8.31; ML: 1.80), (AP: -8.31, ML: 4.00), (AP: -10.5; ML: -1.8) with a reference screw at (AP: -10.5; ML: 2.00).

### Nonlinear Atlas Registration and Anatomical Parcellation

For precise anatomical parcellation of the rodent brain, T2-weighted structural magnetic resonance imaging (MRI) scans from each subject were registered to the Waxholm Space (WHS) Sprague Dawley rat brain atlas (v4.01). We employed a hierarchical, multi-resolution registration pipeline to map the atlas into native subject space. Initially, a global rigid and affine transformation was estimated using SimpleITK, optimizing Mattes Mutual Information via gradient descent to establish a baseline macroscopic alignment. This initialization was followed by a dense, non-linear deformable registration step implemented within the MONAI framework. To account for local structural variations while preserving topological integrity, the non-linear optimization minimized a Local Normalized Cross-Correlation (LNCC) loss function regularized by a bending energy penalty, ensuring smooth, diffeomorphic transformations. The resulting dense displacement fields (DDFs) were inverted to warp the comprehensive WHS anatomical labels directly into each subject’s native MRI space.

### EEG Probe Induced Mechanical Tissue Displacement

Standard image registration pipelines inherently fail to account for the focal, mechanical tissue displacement caused by the physical insertion of acute neural probes. To correct the registered anatomical label maps for these localized physical distortions, we developed a subject-specific trajectory modeling approach. Leveraging the empirical tip and shaft coordinates of the implanted devices, we computationally generated a synthetic 3D cylindrical deformation field. This localized geometric field was mathematically formulated to simulate the radial expansion of brain tissue induced by the physical volume of the probes, explicitly parameterized by the specific inner and outer diameters of the implanted shanks.

Once constructed, these synthetic focal deformations were spatially aligned with the localized in vivo probe trajectories and systematically applied to the registered subject-space atlas labels. Administered via nearest-neighbor interpolation to preserve discrete anatomical boundaries, this secondary warping step retroactively models the outward displacement of tissue enveloping the functional electrode arrays. The culmination of this two-stage pipeline constituting global atlas-to-subject normalization followed by localized, probe-induced deformation yields a customized anatomical segmentation that incorporates deformations due to electrode insertions. This corrected mapping accurately reflects the structural reality of the instrumented brain during recording, thereby ensuring precise and reliable anatomical assignments for the recorded electrophysiological units.

## Supplemental Figures

**Supplemental Figure 2.**
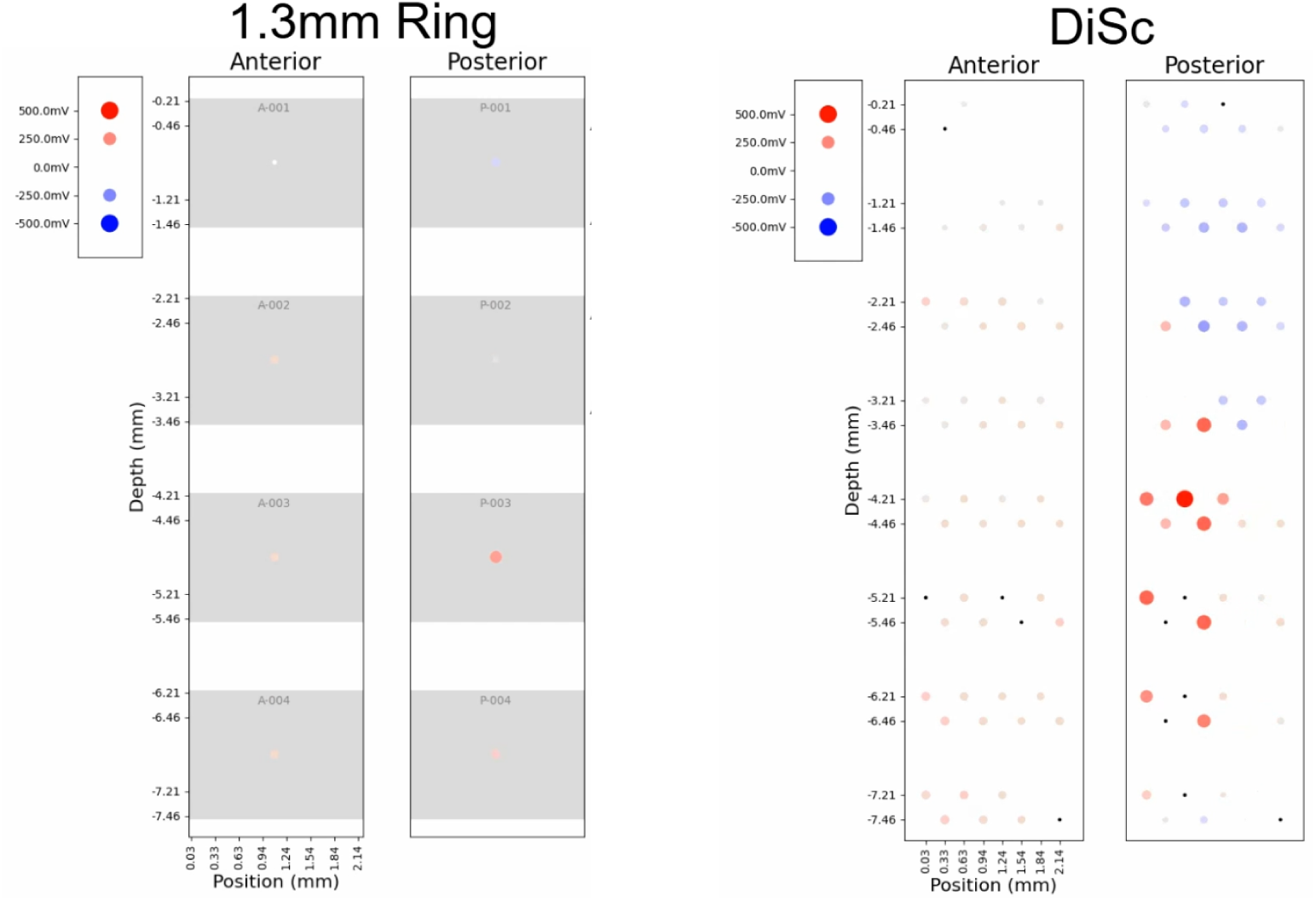
Polarity Visualization. The LFP signals from the two DiSc arrays implanted in Subject 1 are shown as a polarity plot (left), one-second waveform (middle) and five-second waveform (right). In the polarity plot, the voltage at a given electrode is represented as a circle of variable size and color. A large red circle denotes a high-amplitude positive voltage while a large blue circle represents a high-amplitude negative voltage. As the voltage becomes more neutral, the circle shrinks and becomes whiter such that a voltage of zero is invisible. The center plot shows the LFP signal over a short time window to make the dynamics are more easily observed. The right plot shows a wider time window to better express the surrounding activity. Graphs are shown for the data in DiSc configuration (top) and 1.3mm ring configuration (bottom).

**Supplemental Figure 3.**
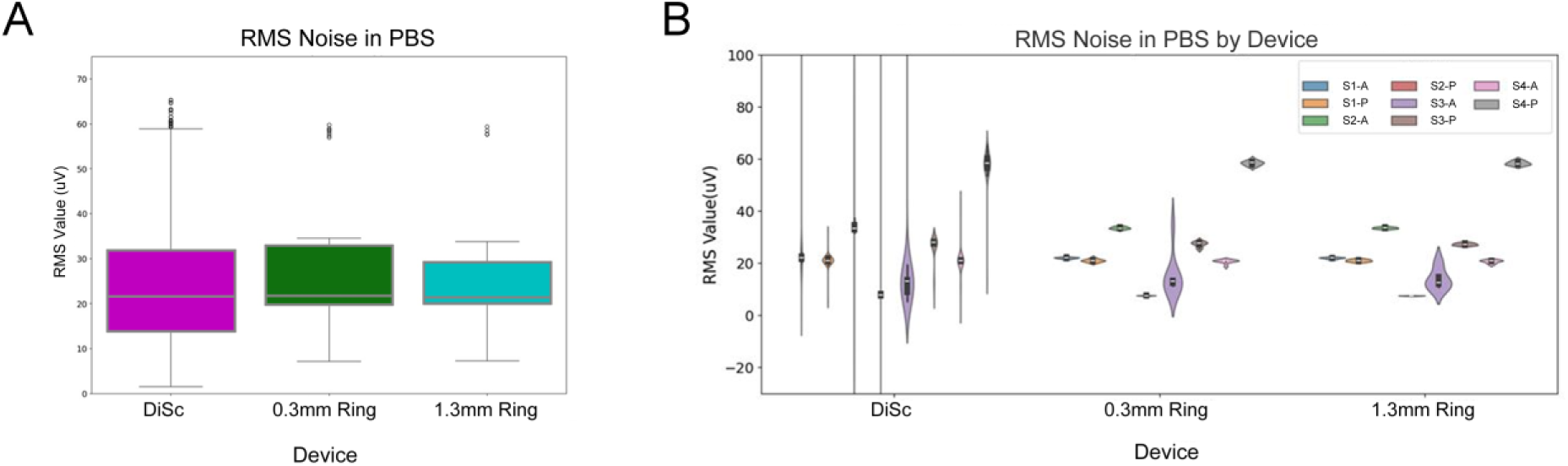
Presurgical RMS Noise. (**A**) Boxplot showing distribution of channel RMS noise values from pre-surgery recordings in phosphate-buffered saline (PBS). Channels are grouped by device configuration. (**B**) Violin plots showing distribution of channel RMS noise values from pre-surgery recordings in PBS. Each violin represents an individual DiSc device. The legend entries denote subject ID – i.e “S1” for Subject 1 – followed by either ‘A’ or ‘P’ if the device was anterior or posterior respectively to the hippocampus.

**Supplemental Figure 4.**
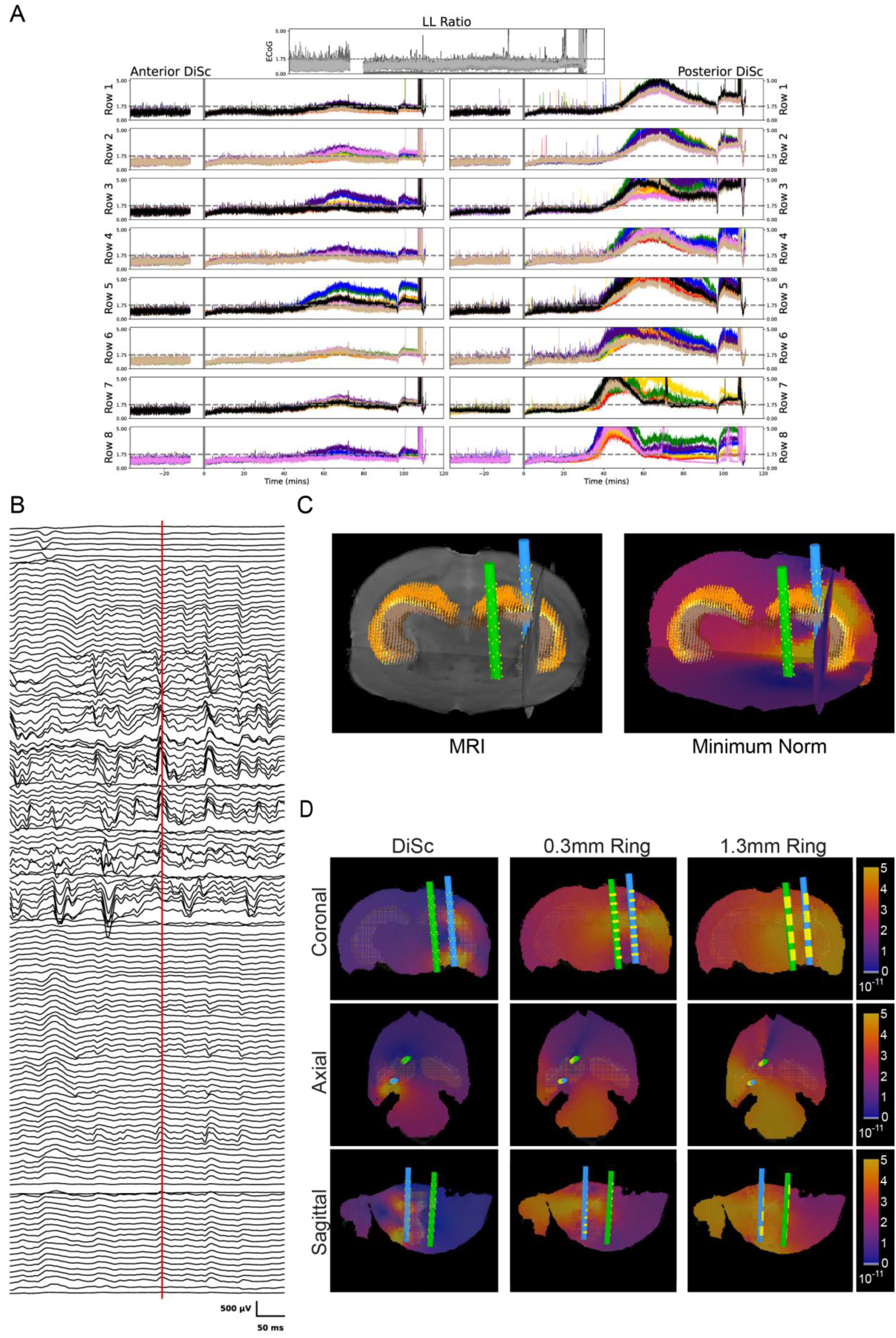
Event from Subject 2 sLORETA. (**A**) Line length ratio over time for Subject 2. Plots use the same conventions as Figure 7-A. (**B**) Excerpt of signal recorded by DiSc arrays in Subject 2. The red line denotes the sample shown in figures C and D. The plot uses the same conventions as Figure 9-A. (**C**) Minimum norm projection of Subject 2 depicting 5015.277s after KA injection. The plot has the same conventions as Figure 9-B. (D) The minimum norm result shown from 3 axes. The plot has the same conventions as Figure 9-C.

**Supplemental Figure 5.**
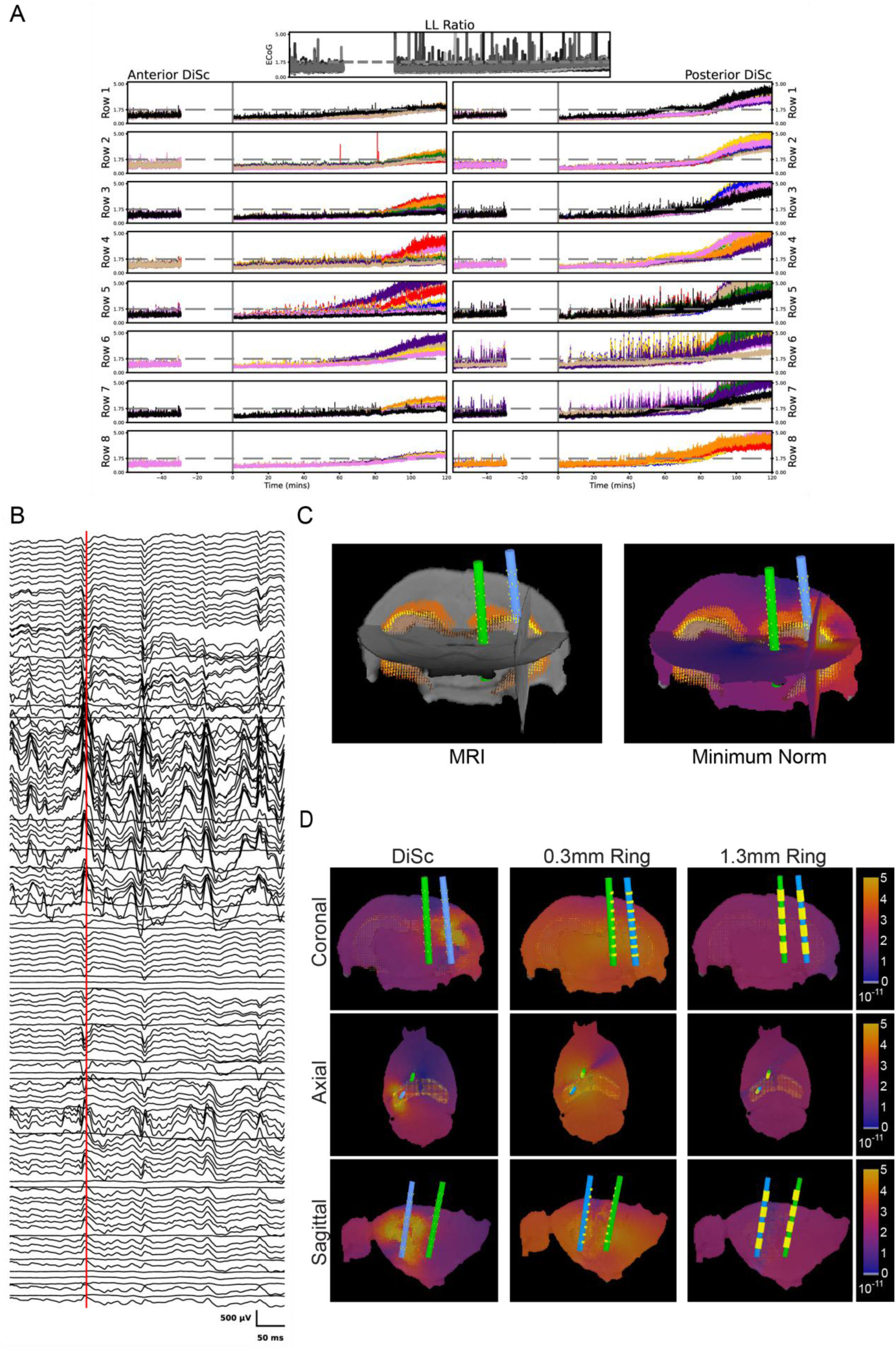
(A) Event from Subject 3 sLORETA. (**A**) Line length ratio over time for Subject 3. Plots use the same conventions as Figure 7-A. (**B**) Excerpt of signal recorded by DiSc arrays in Subject 3. The red line denotes the sample shown in figures C and D. The plot uses the same conventions as Figure 9-A. (**C**) Minimum norm projection of Subject 3 depicting 5921.639s after KA injection. The plot has the same conventions as Figure 9-B. (D) The minimum norm result shown from 3 axes. The plot has the same conventions as Figure 9-C.

**Supplemental Figure 6.**
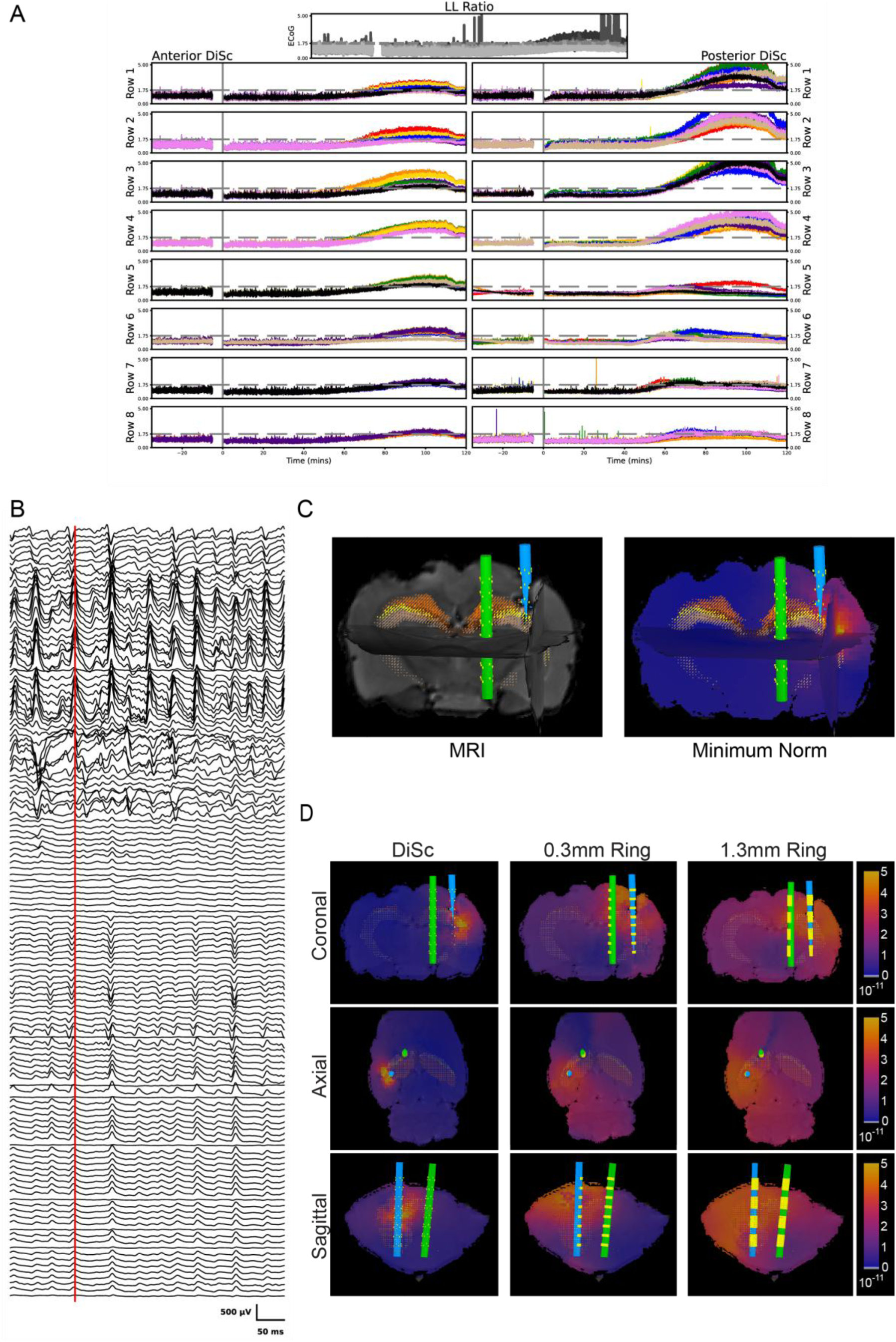
Event from Subject 4 sLORETA. (**A**) Line length ratio over time for Subject 4. Plots use the same conventions as Figure 7-A. (**B**) Excerpt of signal recorded by DiSc arrays in Subject 4. The red line denotes the sample shown in figures C and D. The plot uses the same conventions as Figure 9-A. (**C**) Minimum norm projection of Subject 4 depicting 5728.118s after KA injection. The plot has the same conventions as Figure 9-B. (D) The minimum norm result shown from 3 axes. The plot has the same conventions as Figure 9-C.

**Supplemental Figure 7.**
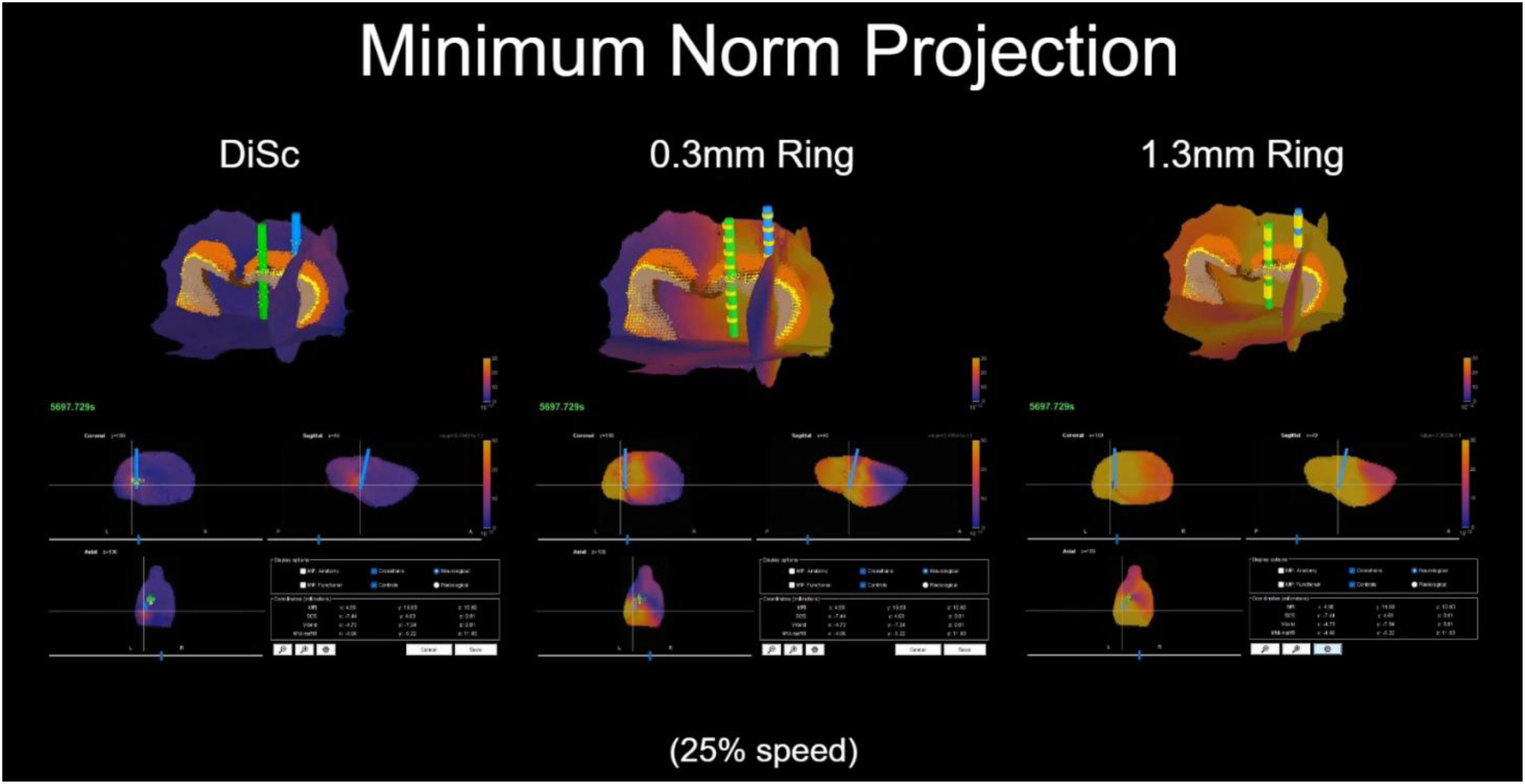
Minimum Norm Projection Video. A video depicting the minimum norm result for Subject 1 from two seconds prior to three seconds following the event shown in Figure 9. The results are shown for each device configuration to show the resolution differences over time. Videos have been slowed to quarter and half speed so that five seconds of data are displayed over twenty and then ten seconds sequentially.

## References

Abrego, A.M. et al. (2023) “Sensing local field potentials with a directional and scalable depth electrode array,” Journal of Neural Engineering, 20(1), p. 016041. Available at: 10.1088/1741-2552/acb230.

Ben-Ari, Y., Tremblay, E. and Ottersen, O.P. (1980) “Injections of kainic acid into the amygdaloid complex of the rat: An electrographic, clinical and histological study in relation to the pathology of epilepsy,” Neuroscience, 5(3), pp. 515–528. Available at: 10.1016/0306-4522(80)90049-4.

Chowdhury, F.A. et al. (2021) “Localisation in focal epilepsy: a practical guide,” Practical Neurology, 21(6), pp. 481–492. Available at: 10.1136/practneurol-2019-002341.

Esteller, R. et al. (2001) “Line length: an efficient feature for seizure onset detection.” *Conference Proceedings of the 23rd Annual International Conference of the IEEE Engineering in Medicine and Biology Society*, Istanbul, Turkey.

Esteller, R., Echauz, J. and Tcheng, T. (2004) “Comparison of line length feature before and after brain electrical stimulation in epileptic patients.” *The 26th Annual International Conference of the IEEE Engineering in Medicine and Biology Society*, San Francisco, CA.

Jayalakshmi, S. et al. (2023) “Long-Term Seizure Freedom, Resolution of Epilepsy and Perceived Life Changes in Drug Resistant Temporal Lobe Epilepsy With Hippocampal Sclerosis: Comparison of Surgical Versus Medical Management,” Neurosurgery, 92(6), pp. 1249–1258. Available at: 10.1227/neu.0000000000002358.

Jehi, L. et al. (2021) “Comparative Effectiveness of Stereotactic Electroencephalography Versus Subdural Grids in Epilepsy Surgery,” Annals of Neurology, 90(6), pp. 927–939. Available at: 10.1002/ana.26238.

Kwan, P. and Brodie, M.J. (2000) “Early identification of refractory epilepsy,” The New England Journal of Medicine, 342(5), pp. 314–319. Available at: 10.1056/NEJM200002033420503.

Medani, T. et al. (2023) “Brainstorm-DUNEuro: An integrated and user-friendly Finite Element Method for modeling electromagnetic brain activity,” NeuroImage, 267, p. 119851. Available at: 10.1016/j.neuroimage.2022.119851.

Medani, T. et al. (2025) “High-Resolution Directional Depth Electrodes: Open-Source FEM Lead-Field Modeling, Characterization, and Validation,” *arXiv preprint* [Preprint].

van Mierlo, P. et al. (2020) “Ictal EEG source localization in focal epilepsy: Review and future perspectives,” Clinical Neurophysiology, 131(11), pp. 2600–2616. Available at: 10.1016/j.clinph.2020.08.001.

Ren, X. et al. (2021) “Connectivity and Neuronal Synchrony during Seizures,” The Journal of Neuroscience, 41(36), pp. 7623–7635.

Rusina, E., Bernard, C. and Williamson, A. (2021) “The Kainic Acid Models of Temporal Lobe Epilepsy,” eNeuro, 8(2), p. ENEURO.0337-20.2021. Available at: 10.1523/ENEURO.0337-20.2021.

Sherman, E.M.S. et al. (2011) “Neuropsychological outcomes after epilepsy surgery: Systematic review and pooled estimates,” Epilepsia, 52(5), pp. 857–869. Available at: 10.1111/j.1528-1167.2011.03022.x.

Stacey, W.C. et al. (2012) “Signal distortion from microelectrodes in clinical EEG acquisition systems,” Journal of Neural Engineering, 9(5). Available at: https://pubmed.ncbi.nlm.nih.gov/22878608/.

Stead, M. et al. (2010) “Microseizures and the spatiotemporal scales of human partial epilepsy,” Brain, 133(9), pp. 2789–2797. Available at: 10.1093/brain/awq190.

Sun, J. et al. (2022) “Intraoperative microseizure detection using a high-density micro- electrocorticography electrode array,” Brain Communications, 4(3), p. fcac122. Available at: 10.1093/braincomms/fcac122.

Tadel, F. et al. (2011) “Brainstorm: A user-friendly application for MEG/EEG analysis,” Computational Intelligence and Neuroscience, 2011, p. 879716. Available at: 10.1155/2011/879716.

Téllez-Zenteno, J.F. and Hernández-Ronquillo, L. (2012) “A review of the epidemiology of temporal lobe epilepsy,” Epilepsy Research and Treatment, 2012, p. 630853. Available at: 10.1155/2012/630853.

Vakharia, V.N. et al. (2018) “Getting the best outcomes from epilepsy surgery,” Annals of Neurology, 83(4), pp. 676–690. Available at: 10.1002/ana.25205.

Wiebe, S. et al. (2001) “A randomized, controlled trial of surgery for temporal-lobe epilepsy,” The New England Journal of Medicine, 345(5), pp. 311–318. Available at: 10.1056/NEJM200108023450501.

Willis, J.A. et al. (2026) “Optimizing electrode placement and information capacity for local field potentials in cortex,” NeuroImage, 327, p. 121747. Available at: 10.1016/j.neuroimage.2026.121747.

World Health Organization (2024) Epilepsy. Available at: https://www.who.int/news-room/fact-sheets/detail/epilepsy.

Wu, S. et al. (2024) “Depth versus surface: A critical review of subdural and depth electrodes in intracranial electroencephalographic studies,” Epilepsia, 65, pp. 1868–1878. Available at: 10.1111/epi.18002.

Yan, H. and Ibrahim, G.M. (2019) “Resective epilepsy surgery involving eloquent cortex in the age of responsive neurostimulation: A value-based decision-making framework,” Epilepsy & Behavior, 99, p. 106479. Available at: 10.1016/j.yebeh.2019.106479.

